# Redox-mediated activation of ATG3 promotes ATG8 lipidation and autophagy progression in Chlamydomonas

**DOI:** 10.1101/2023.01.24.525316

**Authors:** Manuel J. Mallén-Ponce, María Esther Pérez-Pérez

## Abstract

Autophagy is one of the main degradative pathways used by eukaryotic organisms to eliminate useless or damaged intracellular material in order to maintain cellular homeostasis under stress conditions. Mounting evidence indicates a strong interplay between the generation of ROS and the activation of autophagy. Although a tight redox regulation of autophagy has been shown in several organisms including microalgae, the molecular mechanisms underlying this control remain poorly understood. In this study, we have performed an in-depth *in vitro* and *in vivo* redox characterization of ATG3, an E2-activating enzyme involved in ATG8 lipidation and autophagosome formation, from two evolutionary distant unicellular model organisms: the green microalga *Chlamydomonas reinhardtii* and the budding yeast *Saccharomyces cerevisiae*. Our results indicated that ATG3 activity from both organisms is subjected to redox regulation since these proteins require reducing equivalents to transfer ATG8 to the phospholipid phosphatidylethanolamine. We established the catalytic Cys of ATG3 as redox target in algal and yeast proteins, and showed that the oxidoreductase thioredoxin efficiently reduces ATG3. Moreover, *in vivo* studies revealed that the redox state of ATG3 from *Chlamydomonas reinhardtii* undergoes profound changes in the absence of photoprotective carotenoids, a stress condition that activates autophagy in algae.

## INTRODUCTION

Macroautophagy (hereafter autophagy) is a dynamic and highly conserved recycling mechanism by which eukaryotic cells degrade cytoplasmic components, including protein aggregates or dysfunctional organelles in the vacuole (or lysosome). This catabolic pathway is an adaptive response to different stress conditions that allows cells to maintain cellular homeostasis and cope with stress (He and Klionsky, 2009; Mizushima et al., 2011). Although basal autophagy plays a housekeeping function in the absence of stress, this degradative process is up-regulated to eliminate unneeded cellular material under adverse conditions (Signorelli et al., 2019). Autophagy was initially considered as a bulk degradation pathway to provide building blocks and recycle nutrients, but it is well established that intracellular content can be specifically degraded through selective autophagy. Indeed, selective degradation of organelles and complex macromolecules such as mitochondria (mitophagy), chloroplast (chlorophagy) or proteasome (proteaphagy) has been reported (Kanki et al., 2009; Marshall et al., 2015; Galluzzi et al., 2017; Izumi et al., 2017; Gatica et al., 2018; Adriaenssens et al., 2022).

Autophagy is characterized by the *de novo* biogenesis of double-membrane vesicles or autophagosomes, which engulf and deliver cytoplasmic material or cargo to the vacuole for degradation and recycling (Supplemental Figure S1). Thus, autophagosome formation is a morphological hallmark of autophagy and has been widely used to monitor autophagy activation in different organisms, including the model single-celled microalga *Chlamydomonas reinhardtii* (Kirisako et al., 1999; Yoshimoto et al., 2004; Pérez-Pérez et al., 2010).

Autophagy is carried out by more than 40 autophagy-related or ATG proteins, most of which have been first described in yeasts (Tsukada and Ohsumi, 1993; Ichimura et al., 2000; Díaz-Troya et al., 2008; Liu and Bassham, 2012; Shemi et al., 2015). Autophagosome biogenesis is mediated by a set of ATG proteins that constitute the core autophagy machinery (Xie and Klionsky, 2007; Nakatogawa et al., 2009; Mizushima et al., 2011; Nakatogawa, 2020). Among them, ATG8 plays a crucial function in autophagy and is the only known ATG protein that binds tightly to autophagic membranes throughout the whole autophagy process, from the recruitment of ATG proteins to the phagophore assembly site (PAS) to the release of the autophagic body into the vacuole. ATG8 plays a critical role in autophagosome formation and contributes to cargo recognition and autophagosome tethering to the vacuolar membrane (Kirisako et al., 1999; Ichimura et al., 2000; Kirisako et al., 2000).

To perform these functions, ATG8 must first bind to the phospholipid phosphatidylethanolamine (PE) to form the ATG8-PE adduct in a process known as ATG8 lipidation or ATG8 conjugation (Supplemental Figure S1). Binding of ATG8 to PE occurs through a set of sequential and coordinated reactions governed by the ubiquitin-like ATG8 system (Ichimura et al., 2000; Kirisako et al., 2000). First, nascent ATG8 is processed at a highly conserved Gly at its C-terminus by the Cys-protease ATG4. Then, the E1-like enzyme ATG7 activates and transfers processed ATG8 to the catalytic Cys of the E2-like enzyme ATG3. Finally, ATG3 catalyzes the covalent binding of PE to the C-terminal Gly of ATG8 to form the ATG8-PE conjugate (Ichimura et al., 2000; Kirisako et al., 2000). The ATG12-ATG5 conjugate, formed by the ubiquitin-like ATG12 system, acts as an E3-like ligase that potentiates the final step of ATG8 lipidation by rearranging the catalytic site of ATG3 (Hanada et al., 2007; Sakoh-Nakatogawa et al., 2013). The ATG4 protease also has a deconjugating activity that cleaves ATG8-PE and releases free ATG8 from membranes (Kirisako et al., 2000) (Supplemental Figure S1). Therefore, the activity of proteins from the ATG8-conjugating system controls the balance between free and lipidated ATG8, which is essential for autophagosome formation. Consequently, lipidated ATG8 has been widely used as an autophagy marker due to the correlation between the formation of ATG8-PE and the induction of this catabolic process (Klionsky, 2021).

Autophagy has to be tightly regulated since loss of regulation of this catabolic process results in cellular dysfunction, hypersensitivity to stress and metabolic disorders such as cancer and neurodegenerative diseases in humans (Hofius et al., 2009; Levine and Kroemer, 2019; Signorelli et al., 2019; Klionsky et al., 2021). It has been demonstrated that the Target of Rapamycin (TOR) kinase, a master regulator of cell growth, negatively regulates autophagy in multiple organisms, ranging from unicellular yeasts and algae to multicellular plants and animals (Noda and Ohsumi, 1998; Liu and Bassham, 2010; Pérez-Pérez et al., 2010). In addition to TOR, the SnRK1/AMPK/Snf1 kinase activates autophagy in response to energy or nutrient deficiency (Soto-Burgos and Bassham, 2017). On the other side, mounting evidence indicates that the production of reactive oxygen species (ROS) under stress conditions regulates autophagy in different organisms. In Chlamydomonas, ROS participate in the activation of autophagy under oxidative stress generated by H_2_O_2_ or MV (Pérez-Pérez et al., 2010; Pérez-Pérez et al., 2012a), ER stress (Pérez-Pérez et al., 2010; Pérez-Martín et al., 2014), carotenoid depletion (Pérez-Pérez et al., 2012a), chloroplast damage (Heredia-Martínez et al., 2018), high light stress (Pérez-Pérez et al., 2012a; Kuo et al., 2020), or impaired starch biosynthesis (Tran et al., 2019). In close agreement, autophagy is involved in degrading oxidized proteins under oxidative stress conditions in *Arabidopsis thaliana* (Xiong et al., 2007). Moreover, several studies in plants suggest that SnRK2 activates autophagy in response to ROS produced under abiotic and biotic stress conditions by inhibiting TOR kinase (Rosenberger and Chen, 2018; Signorelli et al., 2019). Thus, an interplay between ROS and autophagy has been described in both plants and algae since ROS activates autophagy as a protective mechanism to decrease ROS production (Pérez-Pérez et al., 2012a; Signorelli et al., 2019).

The control of autophagy by redox signals is an emerging research field and the molecular mechanisms underlying this regulation have not been fully elucidated. The Cys-protease ATG4 is the first ATG protein whose activity has been shown to be redox regulated. In mammals, ATG4 is a direct target of H_2_O_2_ and ROS produced in mitochondria serve as a signaling molecule during autophagy induced by nutrient starvation (Scherz-Shouval et al., 2007). Detailed biochemical studies carried out in yeasts and algae demonstrated that ATG4 is subjected to redox post-translational modifications in these organisms (Pérez-Pérez et al., 2014; Pérez-Pérez et al., 2016). These studies also uncovered the basic mechanism for the redox regulation of yeast and algal ATG4 proteins, which involves the formation of a single disulfide bond with a very low redox potential that can be efficiently reduced by thioredoxin. It has also been shown that ATG4 oxidation leads to further ATG4 inactivation through oligomer formation in yeasts and algae (Pérez-Pérez et al., 2014; Pérez-Pérez et al., 2016; Pérez-Pérez et al., 2021). In consonance with the redox regulation of human, yeast and algal ATG4 proteins, reversible inhibition of ATG4 by H_2_O_2_ has been proposed in plants (Woo et al., 2014).

ATG4 does not seem to be the only ATG protein subject to redox regulation. Under oxidative stress, direct oxidation of human ATG3 and ATG7 at the active site thiols prevents lipidation of LC3 (an ATG8 isoform in mammals) and thus autophagy activation (Frudd et al., 2018). Whether redox regulation of human ATG3 is conserved in other organisms is unknown. In this study, we have performed an in-depth biochemical analysis of the ATG3 protein from *Chlamydomonas reinhardtii* and *Saccharomyces cerevisiae*, two evolutionary distant model organisms with highly conserved autophagy machineries. We have investigated the redox regulation of the ATG8-conjugating activity of ATG3 by *in vitro* and *in vivo* approaches. Our data unraveled the molecular mechanism by which redox signals control ATG3 activity to promote ATG8 lipidation and autophagy progression in Chlamydomonas subjected to oxidative damage generated by carotenoid depletion.

## RESULTS

### ATG3 regulation depends on the redox potential

In order to investigate whether the *Chlamydomonas reinhardtii* ATG3 protein (CrATG3) is subjected to redox regulation, we performed a detailed biochemical analysis of the recombinant protein under different reducing and oxidizing conditions. CrATG3 migrated as a 36 kDa protein in the presence of β-mercaptoethanol (βME) but an additional band of ≈70 kDa likely corresponding to dimeric CrATG3 was observed in non-reducing denaturing gels (Supplemental Figures S2A and S2B). Incubation of CrATG3 with reducing agents such as DTT_red_ or GSH promotes CrATG3 monomerization, whereas oxidation with DTT_ox_, H_2_O_2_ or Cu^2+^ led to protein dimerization and the detection of an oxidized form of monomeric CrATG3 (Figure 1A). Moreover, both CrATG3 reduction and oxidation were fully reversible processes, strongly suggesting the involvement of a redox post-translational modification (PTM) in the regulation of this protein (Figure 1B). Accordingly, increasing DTT_red_ concentrations or longer incubations with this reducer boosted CrATG3 monomerization (Figures S2C and S2D) while raising H_2_O_2_ concentrations increased the level of dimeric and oxidized monomeric CrATG3 (Figure S2E).

**Figure 1.**
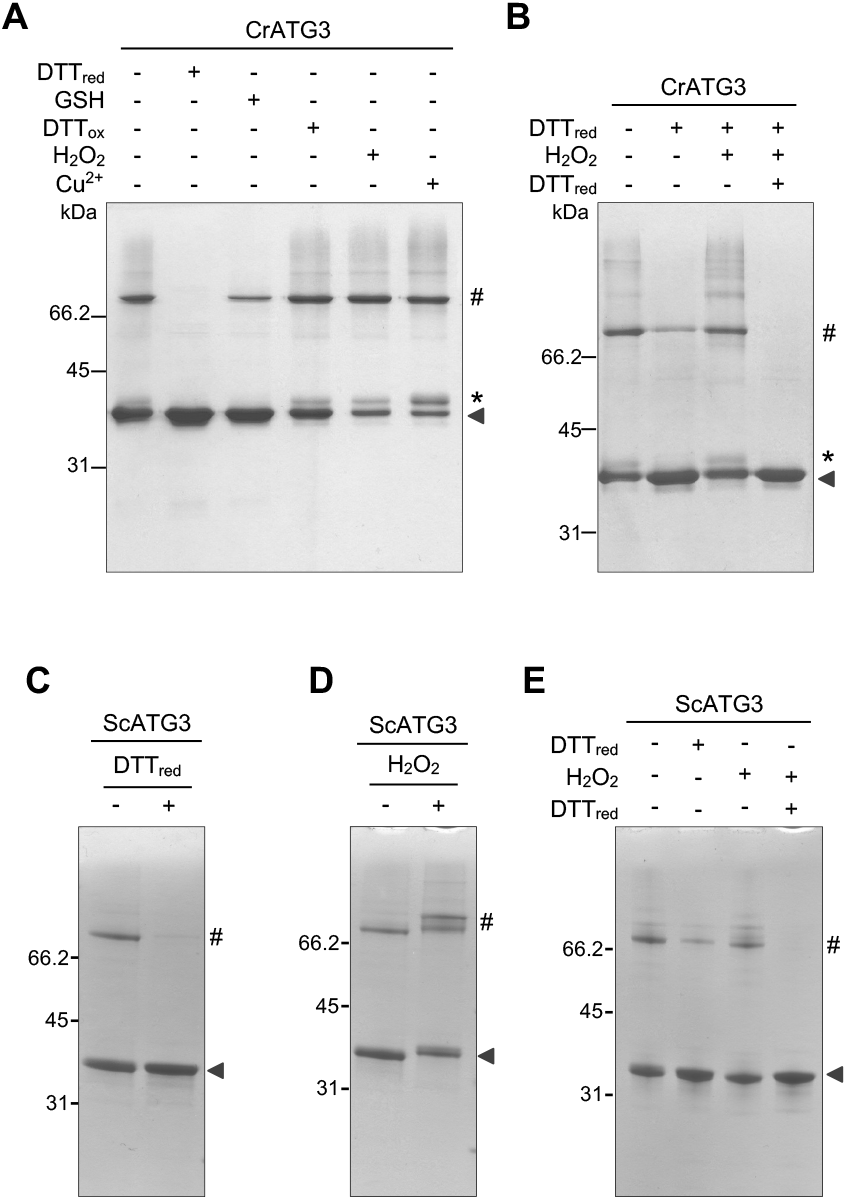
Reversible oxidation of ATG3 from *Chlamydomonas reinhardtii* and *Saccharomyces cerevisiae*. **A.** Chlamydomonas ATG3 (CrATG3) untreated (-) or treated (+) with different redox agents such as DTT_red_ (2.5 mM), GSH (5 mM), DTTox (2.5 mM), H_2_O_2_ (0.5 mM) or Cu^2+^ (0.1 mM) for 30 min and then resolved by 12% non-reducing SDS-PAGE gel. **B.** CrATG3 untreated (-) or treated (+) with the reducer DTT_red_ (0.5 mM, 30 min), then with the oxidant H_2_O_2_ (1 mM, 20 min), and finally with a higher DTT_red_ concentration (10 mM, 30 min). **C.** Saccharomyces ATG3 (ScATG3) untreated (-) or treated (+) with DTT_red_ (2.5 mM, 30 min). **D.** ScATG3 in the absence (-) or presence (+) of H_2_O_2_ (0.5 mM, 30 min). **E.** ScATG3 was sequentially treated (+) with 0.5 mM DTT for 30 min, then with 1 mM H_2_O_2_ for 20 min and finally with 10 mM DTT for 30 min. All incubation assays were performed at 25°C using 2.5 μg of purified CrATG3 (*A* and *B*) or ScATG3 (*C-E*). In all experiments, untreated sample (-) was used as control. The molecular mass marker (kDa) is shown on the left. The arrow indicates reduced monomeric CrATG3 or ScATG3. The symbols * and # indicate an oxidized monomeric CrATG3 and dimeric CrATG3 or ScATG3, respectively.

Next, we investigated whether ATG3 from other organisms is redox regulated. To this aim, we focused on ATG3 from the model yeast *Saccharomyces cerevisiae* (ScATG3) because this organism is evolutionarily distant from Chlamydomonas and the protein shares only 32% identity with CrATG3 (Supplemental Figures S3A and S3B). First, we performed a redox characterization of the recombinant ScATG3 (Supplemental Figures S2F). As described for CrATG3, ScATG3 migrated as monomeric or dimeric forms in non-reducing gels depending on the redox conditions. We found that the presence of reducing agents including βME (Supplemental Figure S2G) or DTT_red_ (Figure 1C) promotes ATG3 monomerization, whereas the addition of H_2_O_2_ resulted in ATG3 dimerization and the detection of higher oligomerized forms of the protein (Figure 1D). Both reduction and oxidation were found to be reversible, strongly suggesting that ScATG3 was also subjected to redox PTM (Figure 1E).

The similar results obtained with two divergent ATG3 enzymes, CrATG3 and ScATG3, prompted us to further characterize their redox regulation. To investigate the link between ATG3 and the redox potential, we performed a complete DTT redox titration by varying the ambient redox potential (*E*_h_ at pH 7.5) between −262 and −374 mV. CrATG3 and ScATG3 proteins were incubated at each redox potential and the ratio of ATG3 monomerization was determined by analyzing the electrophoretic mobility in non-reducing denaturing gels (Figure 2). Our data revealed a strong correlation between the monomerization of both ATG3 proteins and the redox potential. Oxidized (dimer and oxidized monomer) and reduced (monomer) ATG3 forms were observed under less electronegative redox potential (from −262 to −322/-327 mV), although only the reduced monomeric form was detected at *E*_h_ values higher than −345 mV (Figures 2A and 2B). The experimental data of in-gel redox titrations gave good fits to the Nernst equation for the reduction of a single 2-electron component, with a midpoint redox potential (*E*_m_ at pH7.5) of −311.9 mV for CrATG3 (Figure 2C) and −312.9 mV for ScATG3 (Figure 2D), indicating the participation of a redox-regulated disulfide bond in the formation of ATG3 dimers in both proteins. In addition to this modification, CrATG3 showed a clear oxidation of the monomeric form under less electronegative redox potential (Figure 2A) that was not evident for ScATG3 (Figure 2B), suggesting that additional redox PTM may occur *in vivo* in CrATG3.

**Figure 2.**
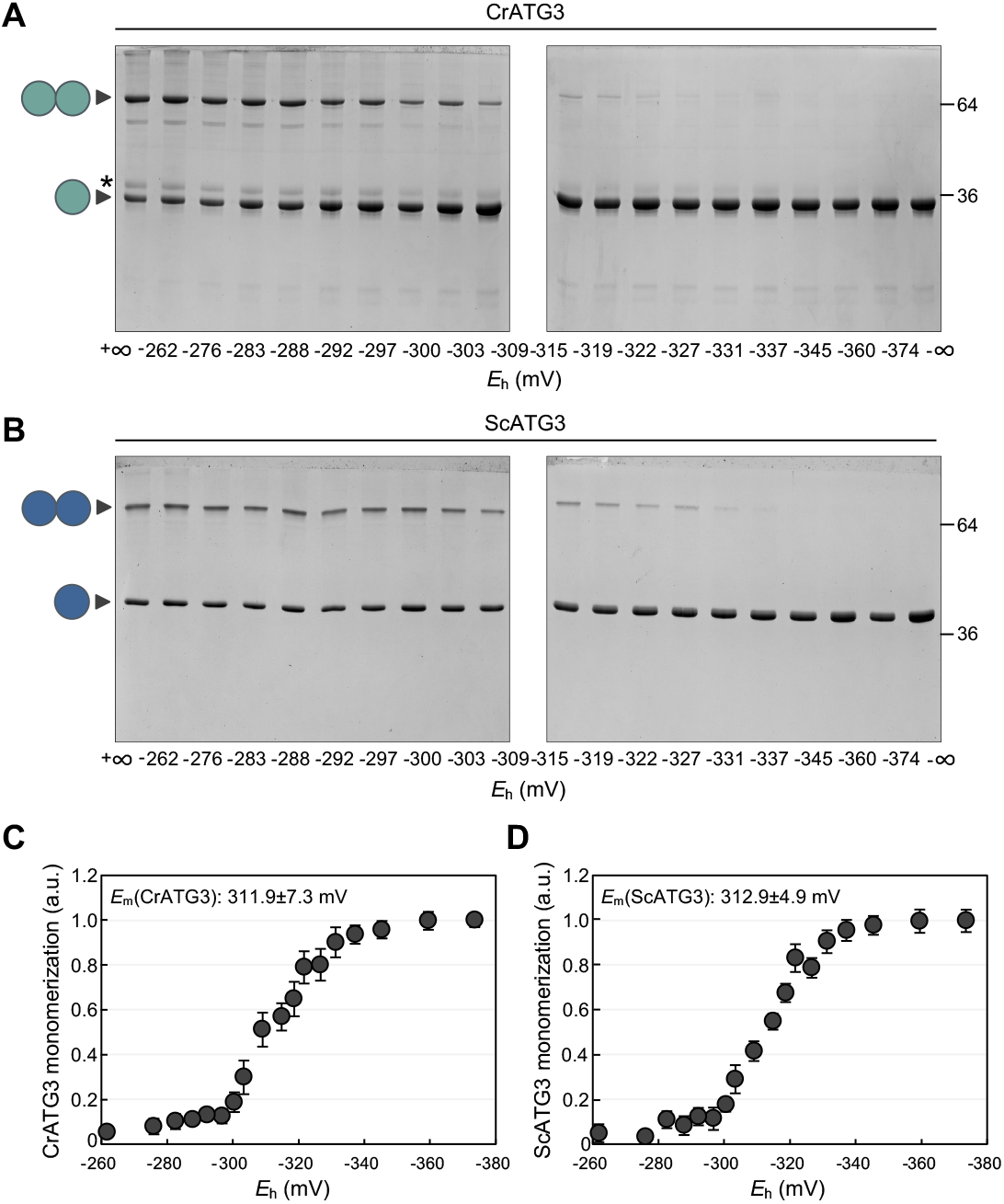
Monomerization of Chlamydomonas and Saccharomyces ATG3 proteins depends on redox potential. **A.** Redox titration of Chlamydomonas ATG3 (CrATG3) monomerization. The different isoforms (dimer, oxidized monomer and reduced monomer) of Chlamydomonas ATG3 were analyzed after incubation at indicated *E*_h_ poised by 20 mM DTT in various dithiol/disulfide ratios. All samples were resolved by non-reducing SDS-PAGE gels and then visualized by Coomassie Brilliant Blue staining. The -∞ sample was considered 100% of CrATG3 reduced monomer and used as reference for quantification. **B.** Redox titration of Saccharomyces ATG3 (ScATG3) monomerization. Analysis of Saccharomyces ATG3 dimer/monomer ratio at the indicated *E*_h_ as described in *(A)*.**C.**-**D.** ATG3 monomerization as monitored in *(A)* and *(B)*, respectively, was quantified and interpolated by nonlinear regression of the data using the Nernst equation for 2 electrons exchanged (n=2) and one redox component. The average midpoint redox potential (*E*_m_,7.5) of 3 independent experiments is shown in the figure as mean ± standard deviation. The symbol * shows oxidized and monomeric CrATG3. One ball and two balls correspond to reduced monomer and dimer, respectively, of CrATG3 or ScATG3.

Thus, taking together our results indicated that CrATG3 and ScATG3 are redox regulated. Moreover, the molecular mechanism of ATG3 redox regulation is likely conserved between yeasts and algae and involves a dithiol-disulfide exchange reaction with a very low midpoint redox potential.

### Thioredoxin efficiently reduces CrATG3

Thioredoxins (Trx) are small oxidoreductases able to reduce disulfide bonds with very low redox potential in target proteins involved in a wide range of cellular processes such as the Calvin-Benson-Bassham (CBB) cycle or stress response (Buchanan and Balmer, 2005; Pérez-Pérez et al., 2017). We have previously demonstrated that Trx plays a role in autophagy by regulating the ATG4 protease in yeasts and algae (Pérez-Pérez et al., 2014; Pérez-Pérez et al., 2016; Pérez-Pérez et al., 2021). Since our results indicated that ATG3 is subjected to redox modifications (Figures 1 and 2 Supplemental Figure S2), we investigated whether Trx may also regulate this essential autophagy protein. To this aim, we tested whether Trx is able to reduce the oxidized forms of CrATG3 by analyzing the time-course monomerization of CrATG3 using Trx and/or DTT_red_ as electron donors. Specifically, we incubated Chlamydomonas TRXh1 (CrTRXh1), the major cytosolic Trx (Lemaire and Miginiac-Maslow, 2004), with CrATG3 for 5, 15, 30 and 60 min and then analyzed the electrophoretic mobility of CrATG3 in non-reducing denaturing gels. Our data showed that CrTRXh1 efficiently reduces and monomerizes CrATG3 in the presence of 0.25 mM DTT_red_ (Figure 3A), which is required to keep Trx reduced (Figure 3B) (Buchanan and Balmer, 2005). However, no effect on CrATG3 monomerization was detected in the absence of CrTRXh1 or when only DTT_red_ was added (Figure 3A). Moreover, the band corresponding to oxidized monomeric CrATG3 also decreased when samples were incubated with CrTRXh1 (Figure 3A). Therefore, our findings indicated that Trx efficiently reduces CrATG3, suggesting a function of this oxidoreductase in the regulation of the redox state of ATG3.

**Figure 3.**
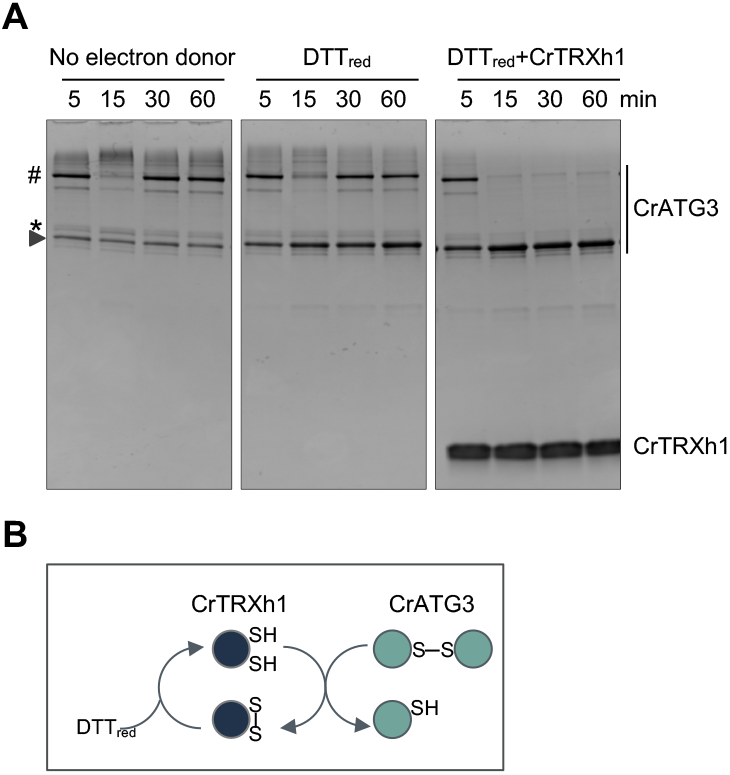
Chlamydomonas ATG3 is reduced by thioredoxin. **A.** Monomerization of CrATG3 after incubation for the indicated times (5, 15, 30, 60 min) in the presence of CrTRXh1 (5 μM) (right panel). A low DTT_red_ concentration (0.25 mM) was used to keep CrTRXh1 at its reduced status. Samples with no reducer (left panel) or only with a low DTT_red_ concentration (middle panel) were used as controls. Proteins were subjected to non-reducing 15% SDS-PAGE gels and then visualized by Coomassie Brilliant Blue staining. The different CrATG3 isoforms and CrTRXh1 are indicated on the right. The reduced and monomeric CrATG3, the oxidized and monomeric CrATG3 and the dimeric CrATG3 are highlighted with an arrow, an asterisk (*) and a hash (#), respectively, on the left. **B.** Schematic representation of the proposed mechanism of monomerization of CrATG3 by the oxidoreductase CrTRXh1. Reduced CrTRXh1 (SH) is able to reduce the disulfide bond of dimeric CrATG3 resulting in reduced and monomerized CrATG3 (SH). DTT_red_ keeps CrTRXh1 in its reduced form. S–S represents a disulfide bond.

### The catalytic Cys of ATG3 is involved in the redox regulation of ATG3

The amino acid sequence of CrATG3 contains three Cys residues: the catalytic Cys (Cys255) and two N-terminal Cys (Cys50 and Cys81) (Figures 4A and Supplemental Figure S4). In order to identify the redox-sensitive Cys(s) of CrATG3, we analyzed the electrophoretic mobility of WT and Cys-to-Ser mutant versions of CrATG3 under different redox conditions in non-reducing denaturing gels (Figure 4). Our data showed that mutation of catalytic Cys255 fully prevented the formation of dimeric CrATG3 regardless of the redox conditions (Figures 4B and 4C). On the contrary, the CrATG3^C50S^ and CrATG3^C81S^ mutants displayed the same behavior as CrATG3^WT^ when treated with DTT_red_ or H_2_O_2_, indicating that these two Cys do not participate in CrATG3 dimerization (Figures 4B and 4C). Moreover, as shown for CrATG3^WT^ (Figure 1B), both reduction and oxidation of CrATG3^C50S^ and CrATG3^C81S^ are fully reversible when proteins are sequentially treated with 0.5 mM DTT_red_, 2.5 mM H_2_O_2_ and then 10 mM DTT_red_ (Figure 4D). We also analyzed whether Cys255 is involved in the oxidation of monomeric CrATG3 by performing a DTT redox titration of this mutant form. Our results indicated that CrATG3^C255S^ is monomeric independently of the redox potential and, interestingly, the oxidized monomeric CrATG3 form was not detected even under the lowest electronegative redox potential (Supplemental Figure S5).

**Figure 4.**
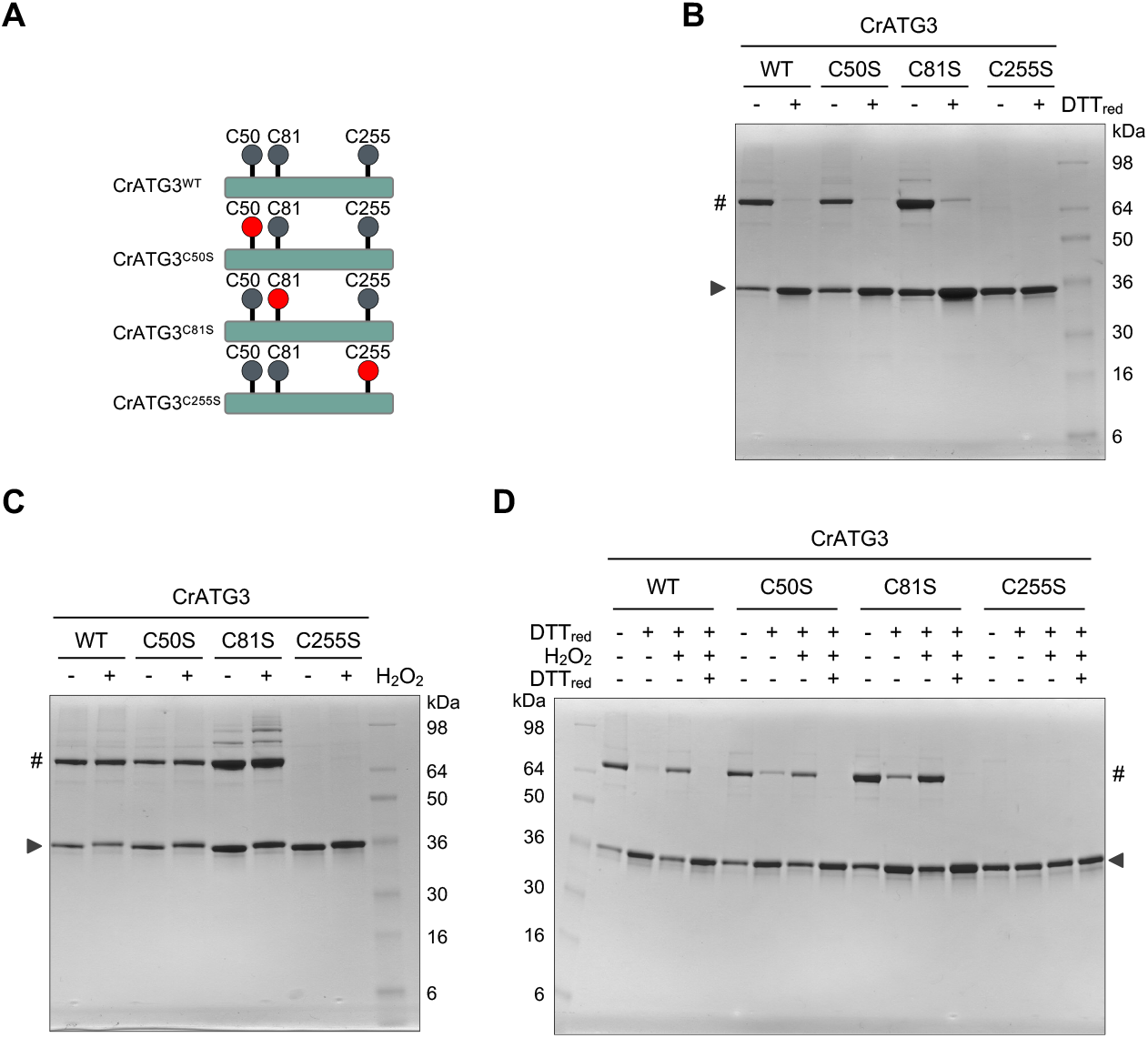
The catalytic Cys of Chlamydomonas ATG3 is subjected to redox regulation. **A.** Schematic representation of WT and Cys-to-Ser mutant versions of CrATG3: CrATG3^WT^, CrATG3^C50S^, CrATG3^C81S^ and CrATG3^C255S^. All Cys residues present in each CrATG3 version are depicted as grey balls, and the Cys-to-Ser mutation is highlighted as a red ball. **B.** Electrophoretic mobility of CrATG3^WT^ (WT), CrATG3^C50S^ (C50S), CrATG3^C81S^ (C81S) and CrATG3^C255S^ (C255S) in the absence (-) or presence (+) of 2.5 mM DTT_red_ for 30 min and then resolved by Coomassie Briliant blue-stained non-reducing SDS-PAGE gels. **C.** All versions of CrATG3 untreated (-) or treated (+) with 0.5 mM H_2_O_2_ for 20 min. **D.** Each CrATG3 version was sequentially incubated (+) with 0.5 mM DTT for 30 min, then with 2.5 mM H_2_O_2_ for 20 min and newly with 10 mM DTT for 45 min. All incubation assays were performed at 25°C using 2.5 μg of each purified Chlamydomonas ATG3 version. In all experiments, untreated sample (-) was used as control. The arrow indicates monomeric CrATG3 and # symbol CrATG3 dimer, respectively. The molecular mass marker (kDa) is shown.

In close agreement with our experimental results, modelling of CrATG3 and ScATG3 proteins predicted that the catalytic Cys is the only highly conserved Cys in these proteins (Supplemental Figures S4B-D). Therefore, we also investigated the role of the catalytic Cys from ScATG3 (Cys234) in the redox regulation of this protein by analyzing the electrophoretic mobility of the Cys-to-Ser catalytic mutant ScATG3^C234S^ (Supplemental Figure S2H). Our results showed that mutation of Cys234 precluded the dimerization of ScATG3 (Supplemental Figure S2H). Thus, taken together, our data strongly suggested that the catalytic Cys of ATG3 is subjected to redox regulation in algae and yeasts.

### ATG3 must be reduced to perform ATG8 lipidation

ATG3 is an E2-activating enzyme that catalyzes the conjugation of ATG8 from ATG7 to the headgroup of PE (Ichimura et al., 2000) (Supplementary Figure S1). To study the implications of CrATG3 redox regulation in the ATG8 conjugation process, we first set up a cell-free lipidation assay using total extract from Chlamydomonas and recombinant CrATG3 protein (Figure 5A). This assay is based on a previous one performed to monitor ATG4 activity in total extracts from Chlamydomonas (Pérez-Pérez et al., 2010; Pérez-Pérez et al., 2016; Crespo and Pérez-Pérez, 2022) and Arabidopsis (Laureano-Marín et al., 2020). ATG8 lipidation can be easily monitored *in vivo* in Chlamydomonas cells subjected to autophagy-activating conditions since processed and free ATG8 can be clearly distinguished from the PE-conjugated ATG8 form (Pérez-Pérez et al., 2010) using 15% resolving gel without urea (Pérez-Pérez et al., 2010; Crespo and Pérez-Pérez, 2022). Briefly, the cell-free lipidation assay included the following steps: (1) incubation of total extract with recombinant CrATG3, (2) protein electrophoresis under non-reducing denaturing conditions, and (3) immunoblot analysis with anti-CrATG8 antibodies. First, we analyzed CrATG8 lipidation from total extracts in the absence or presence of exogenous recombinant CrATG3^WT^. We found that incubation of Chlamydomonas total extracts with DTT_red_ promotes ATG8-PE formation and, notably, addition of exogenous CrATG3^WT^ further increased the level of lipidated CrATG8 (Figure 5B and Supplemental Figure S6A). Next, we monitored ATG8 lipidation in total extracts incubated with CrATG3^WT^ under reducing (DTT_red_) or oxidizing (H_2_O_2_) conditions. While addition of DTT_red_ triggered ATG8-PE formation, no lipidation was detected under non-reducing or oxidizing conditions (Figure 5C). To study the role of each Cys from CrATG3 in ATG8 lipidation, we performed the cell-free assay with the different forms of CrATG3 (CrATG3^WT^, CrATG3^C50S^, CrATG3^C81S^ and CrATG3^C255S^) using DTT_red_ as electron donor. We found that the addition of CrATG3^WT^, CrATG3^C50S^ or CrATG3^C81S^ increases the conjugation of ATG8 to PE whereas the catalytic Cys mutant CrATG3^C255S^ had no effect on ATG8 lipidation (Figure 5D). These results indicated that the catalytic Cys is the only redox-sensitive Cys in CrATG3.

**Figure 5.**
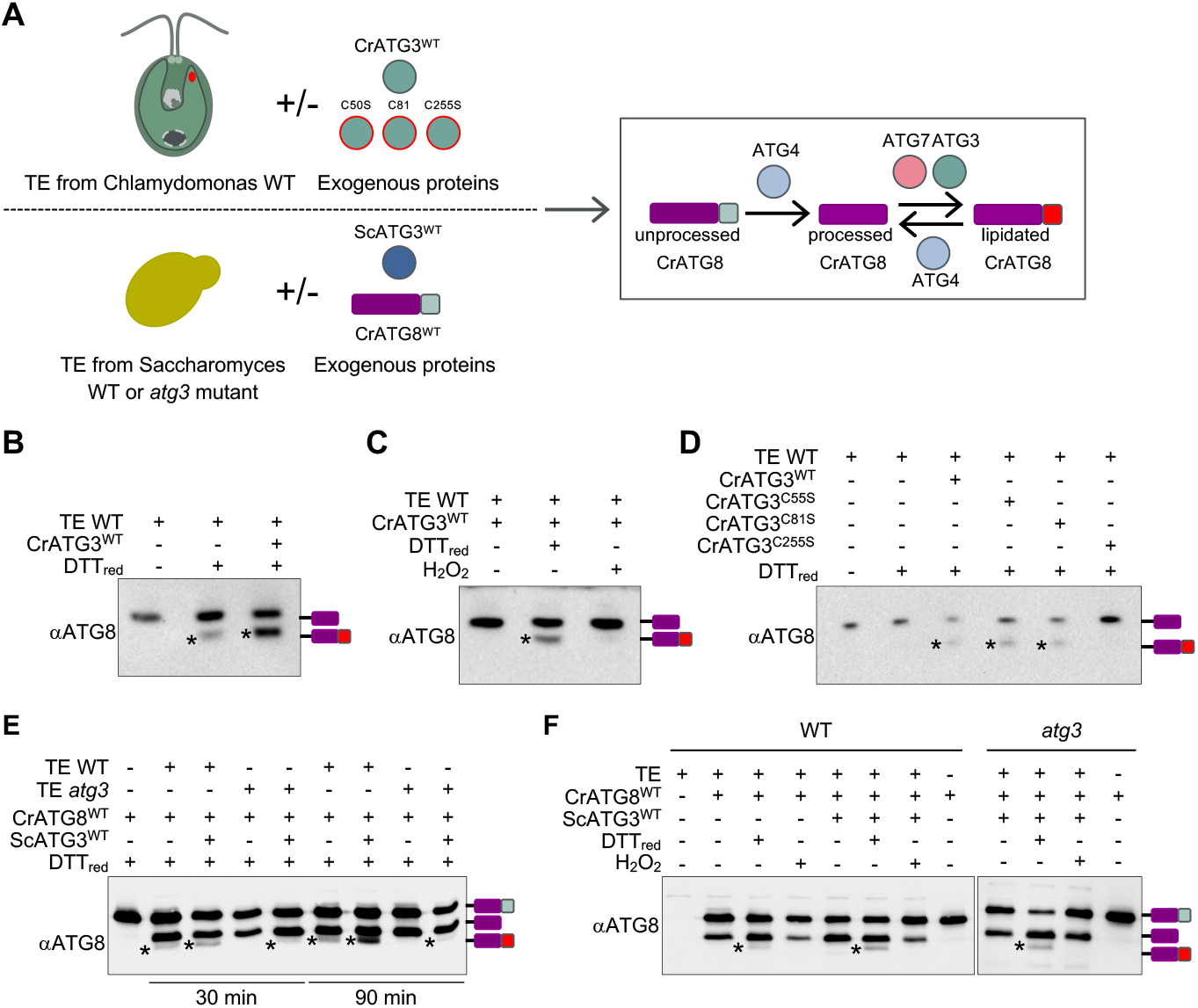
Reduced ATG3 mediates ATG8 lipidation in cell-free assays in Chlamydomonas and Saccharomyces. **A.** Schematic representation of the workflow of the cell-free CrATG8 lipidation assay using the recombinant E2-activating enzyme ATG3 (CrATG3^WT^, CrATG3^C50S^, CrATG3^C81S^, CrATG3^C255S^ or ScATG3^WT^), recombinant CrATG8 (CrATG8^WT^), and total extracts from Chlamydomonas WT or Saccharomyces WT and *atg3* mutant. **B.** Total extracts (TE) from Chlamydomonas WT were untreated (-) or treated (+) with DTT_red_ in the absence (-) or presence (+) of recombinant CrATG3^WT^ for 60 min at 25°C. **C.** Total extracts from Chlamydomonas WT were incubated with no agent (-), a reducer (DTT_red_) or an oxidant (H_2_O_2_) in the presence (+) of recombinant CrATG3^WT^ for 60 min at 25°C. **D.** Total extracts from Chlamydomonas WT were untreated (-) or treated (+) with DTT_red_ in the presence (+) of recombinant CrATG3^WT^, CrATG3^C50S^, CrATG3^C81S^ or CrATG3^C255S^ for 60 min at 25°C. **E.** Total extracts from Saccharomyces WT or *atg3* mutant strains were incubated in a DTT_red_-containing buffer in the absence (-) or presence (+) of recombinant his-tagged ScATG3 (ScATG3^WT^) for 30 or 90 min. **F.** Total extracts from Saccharomyces WT or *atg3* mutant strains were untreated (-) or treated (+) with DTT_red_ or H_2_O_2_ in the absence (-) or presence (+) of recombinant ScATG3^WT^ for 60 min at 25°C. After the lipidation assay, proteins were resolved by 15% SDS-PAGE gel and finally analyzed by western-blot with Chlamydomonas anti-ATG8 antibody (αATG8). Lipidated ATG8 (ATG8-PE) is shown with an asterisk on the gel. The different ATG8 isoforms (unprocessed, processed and lipidated ATG8) are depicted on the right.

Next, we investigated whether the redox regulation of yeast ATG3 might have a role in ATG8 lipidation using a similar cell-free assay with yeast total extracts and recombinant ScATG3. In this assay, recombinant his-tagged CrATG8 protein was used as substrate and its lipidation state was monitored by western-blot with anti-CrATG8 antibodies (Figure 5A). Given that anti-CrATG8 from Chlamydomonas poorly recognizes yeast ATG8 (Pérez-Pérez et al., 2010), no crosstalk signal was detected with endogenous yeast ATG8. To validate our assay, we incubated total extracts from WT and *atg3* yeast strains with CrATG8 under reducing conditions (DTT_red_) in the absence or presence of recombinant ScATG3. As previously reported (Pérez-Pérez et al., 2010), recombinant CrATG8 was efficiently processed by endogenous ATG4 present in yeast total extracts (Figure 5E). In this assay, an additional band of higher mobility was detected when using total extracts from WT yeast cells. We confirmed this band corresponds to lipidated CrATG8 because it was not detected with total extracts from *atg3* mutant cells, which are unable to lipidate ATG8 (Tsukada and Ohsumi, 1993; Ichimura et al., 2000). Our results indicated that endogenous yeast ATG3 lipidates CrATG8, and this lipidation was enhanced when exogenous ScATG3 is added to the reaction mix (Figure 5E and Supplemental Figure S6B). Moreover, addition of exogenous ScATG3 allowed the lipidation of CrATG8 when total extract from *atg3* cells was used (Figure 5E). A time-course assay indicated that ATG8 lipidation increases along time and is potentiated by ScATG3 addition (Figure 5E). Once validated, the cell-free assay allowed us to study the effect of the redox regulation of yeast ATG3 in ATG8 lipidation. In close agreement with the results obtained with CrATG3 (Figure 5C), CrATG8-PE was detected under reducing conditions (DTT_red_) but not upon non-reducing (no treatment) or oxidizing (H_2_O_2_) conditions in total extracts from WT cells (Figure 5F). As expected, CrATG8-PE was not observed in the *atg3* mutant unless ScATG3^WT^ is included in the assay (Figure 5E). Taken together, our findings strongly suggested that the ATG3-mediated-lipidation of ATG8 is regulated by redox signals in algae and yeasts.

### ATG3 is reduced to ensure ATG8 lipidation under autophagy-activating conditions

Our findings about the redox regulation of ATG3 *in vitro* and in cell-free extracts led us to investigate whether this protein is subjected to redox control *in vivo* under ROS-linked autophagy-activating conditions. To this aim, we focused on Chlamydomonas as we have previously demonstrated a clear regulation of autophagy by redox signals in this model organism. We have shown that addition of norflurazon (NF), an herbicide that blocks carotenoid biosynthesis (Sandmann and Albrecht, 1990), to Chlamydomonas cells leads to enhanced ROS production, ATG8 lipidation and ATG4 inactivation, thus resulting in a strong induction of autophagy by redox unbalance (Pérez-Pérez et al., 2012a; Pérez-Pérez et al., 2016).

To investigate the *in vivo* redox regulation of CrATG3, we generated an antibody against this protein (see Materials and Methods) that allowed us to monitor the abundance and redox state of CrATG3 in Chlamydomonas cells under different stress conditions. First, we check that this anti-CrATG3 recognizes a 36 kDa band corresponding to ATG3 protein (Supplemental Figure S7). Then, using standard reducing gels and western blot analysis, we found that CrATG3 abundance increased in NF-treated cells (Figure 6A), in close agreement with the upregulation of CrATG8 and autophagy detected under this stress (Figure 6A) (Pérez-Pérez et al., 2012a). The level of CrATG3 also increased in response to other autophagy-activating conditions such as treatment with cerulenin, a specific inhibitor of fatty acid synthase, or methyl viologen, which generates ROS (Pérez-Pérez et al., 2012a; Heredia-Martínez et al., 2018) (Supplemental Figure S7). Together, these findings indicated that CrATG3 protein abundance increases under ROS-linked stress in Chlamydomonas cells. Next, we examined the redox state of CrATG3 in Chlamydomonas cells upon autophagy activation using an *in vivo* alkylating assay with the blocking agents NEM and MM(PEG)24. NEM binds to free sulfhydryl groups and causes a negligible increase in the molecular weight of the protein (Yano, 2003). The Cys-labeling by NEM occurs only in reduced Cys while MM(PEG)24 binds to originally oxidized Cys that were previously reduced by DTT_red_ in our assay (Figure 6B) (Roberts et al., 2002; Yano, 2003). MM(PEG)_24_ treatment caused an increase in the size of the protein (2.4 kDa) that can be detected by a shift in the mobility of the labeled protein (Figure 6B). Using this *in vivo* alkylating approach, we were able to detect oxidized and reduced forms of CrATG3 in total extracts from Chlamydomonas. Our results indicated that similar amounts of oxidized and reduced CrATG3 were detected in cells under optimal growth conditions (Figure 6C). We then investigated whether NF treatment might influence the redox state of CrATG3. The *in vivo* alkylating analysis revealed profound changes in the redox state of CrATG3 in response to NF treatment. Under normal growth or before autophagy activation, around 60% of total CrATG3 was reduced (Figure 6C). However, induction of autophagy by NF, which was detected around 24 h after treatment, led first to a pronounced oxidation of CrATG3 and then to a complete reduction of the protein at the time of the highest CrATG8 lipidation (48 h) (Figure 6C). Therefore, our study indicated that the *in vivo* redox state of CrATG3 changes in response to oxidative stress to promote CrATG8 lipidation and autophagy progression in NF-treated cells.

**Figure 6.**
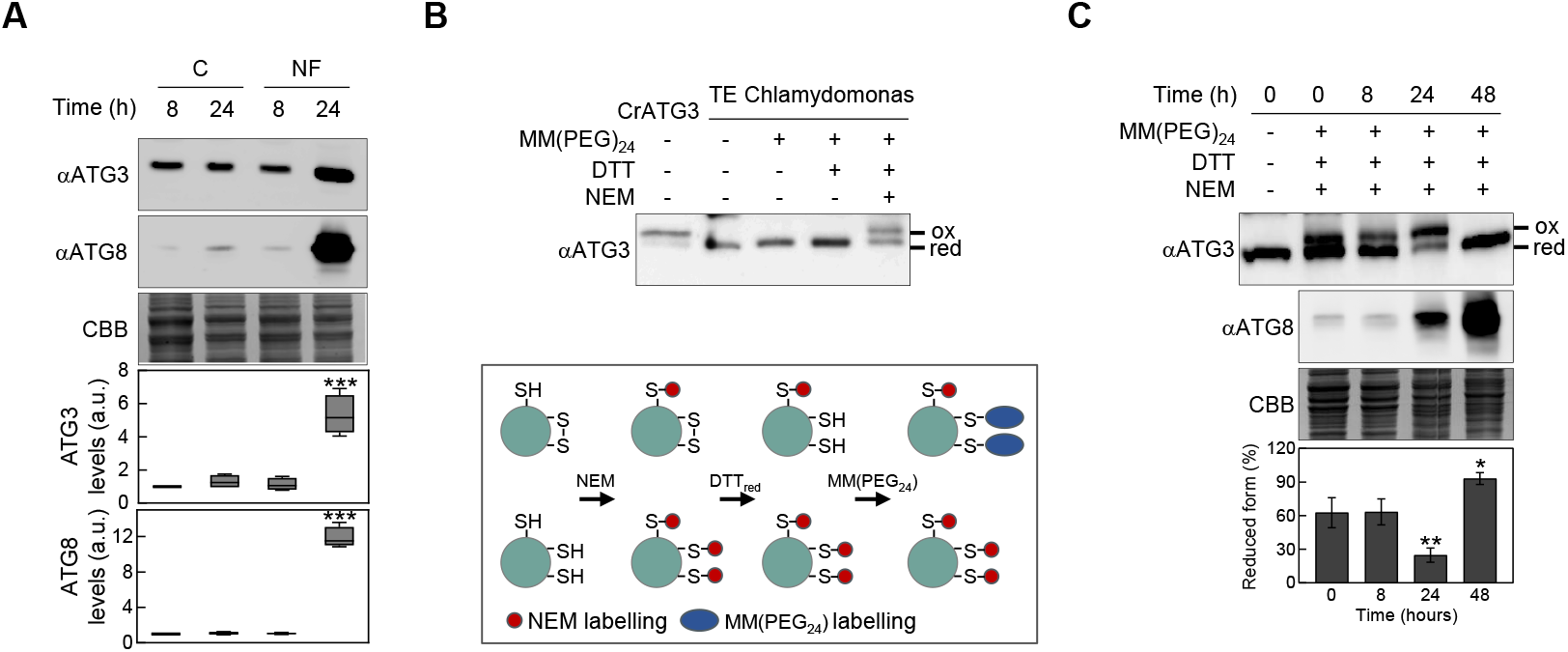
Autophagy-activating conditions leads to a significant increase of ATG3 abundance but also to dynamic changes of ATG3 redox state in Chlamydomonas. **A.** Immunoblot analysis of Chlamydomonas ATG3 (upper panel) and ATG8 (middle panel) in the absence of carotenoids by treatment with 20 μM norflurazon (NF). Untreated cells (C) were used as control. Coomassie Brilliant Blue-stained (CBB) gels were used as protein loading control. Quantification of ATG3 levels (upper box plot) and ATG8 levels (lower box plot) from at least four biological replicates are shown. Asterisks represent significant differences according to two-tailed Student’s t test, ***P < 0.001. **B.** *In vivo* redox state of ATG3 in Chlamydomonas. Proteins were extracted from cells growing exponentially in TAP medium. Then, samples were untreated (-) or treated (+) sequentially with the alkylating (NEM), reducing (DTT_red_) and labeling-alkylating (MM(PEG)24) agents as detailed in Materials and Methods Section. Finally, Chlamydomonas ATG3 was visualized by western blot with anti-CrATG3 (upper panel). His-tagged recombinant CrATG3 (first lane) was used as molecular weight control. CBB gels (lower panel) were used as protein loading control. On the bottom, a schematic representation of the *in vivo* redox alkylating assay is shown. Brieftly: 1) Sulfhydryl groups (SH) are blocked with NEM (red ball). 2) Disulfide bonds (S-S) are reduced by DTT_red_. 3) New SH groups are labeled with MM(PEG_24_) (blue ball). **C.** *In vivo* redox state of ATG3 in NF-treated cells. Proteins were obtained from cells before (0 h) and after (8, 24 and 48 h) NF treatment and labeled with the alkylating agent MM(PEG)24 as described in (*B*). Immunoblotting with anti-CrATG3 (upper panel) and anti-CrATG8 (lower panel). red: reduced form; ox: oxidized form. CBB gels were used as protein loading control. Quantification of reduced ATG3 (%) from four independent experiments is shown. Asterisks represent significant differences according to one-way ANOVA and Bonferroni’s test, **P < 0.01 and *P < 0.05.

## DISCUSSION

Genomic and functional studies have demonstrated that the central machinery of autophagy is conserved in plants and microalgae, but the underlying molecular mechanisms regulating this catabolic process are not fully understood in the green lineage. In this study, we have shown that ATG3, a key autophagy protein required for ATG8 lipidation, is regulated through dynamic reduction/oxidation reactions according to the redox potential in order to ensure autophagy progression under ROS-linked stress in Chlamydomonas.

The maintenance of intracellular redox balance is critical for all organisms, and in particular for photosynthetic organisms since electron transport in the chloroplast generates ROS that can oxidize cellular components including proteins, lipids or nucleic acids (Li et al., 2009)). Photosynthetic cells have multiple layers of defense against oxidative stress, including antioxidant compounds such as glutathione, ascorbate or carotenoids, and antioxidant enzymes such as thioredoxins, catalase or peroxidases (Noctor et al., 2018). Other mechanisms contributing to maintain redox homeostasis in eukaryotic photosynthetic cells involve degradation pathways such as autophagy, which copes with ROS-driven cellular damage to maintain cellular homeostasis (Pérez-Pérez et al., 2012b; Signorelli et al., 2019). In addition to the deleterious effect of ROS, these molecules also act as important signals that mediate the response to stress (Apel and Hirt, 2004; Zaffagnini et al., 2018). Indeed, many proteins can be regulated by the redox state of key Cys residues, including the formation of disulfide bonds, sulfenic acid, or other redox PTMs such as glutathionylation or nitrosylation (Buchanan, 1991; Michelet et al., 2013; Pérez-Pérez et al., 2017; Wakao and Niyogi, 2021). These redox modifications are the result of internal and/or external stimuli that will finally determine subcellular location, protein structure, protein function or protein-protein interactions to allow cells to react and adapt to different environmental cues (Mittler et al., 2011; Choudhury et al., 2017). Importantly, these Cys modifications are fully reversible, resulting in fine protein modulation in response to slight changes of intracellular ROS levels (Zaffagnini et al., 2018; Wakao and Niyogi, 2021).

In order to tackle the harmful effects of ROS, cells up-regulate autophagy to eliminate impaired components including oxidized proteins, protein complexes or even entire organelles such as mitochondria or chloroplast whose dysfunctional activity might further increase ROS levels (Pérez-Pérez et al., 2012b; Signorelli et al., 2019). ROS also act as secondary messengers to modulate the activity of autophagy-related proteins (Scherz-Shouval et al., 2007). The ATG4 protease was the first ATG protein shown to be redox regulated. A pioneer study in mammals demonstrated that HsATG4A and HsATG4B are regulated by direct oxidation of Cys81, an adjacent Cys of the catalytic Cys77 (Scherz-Shouval et al., 2007). However, this regulatory Cys from ATG4 is not widely conserved, suggesting that alternative mechanisms may operate in other organisms to regulate ATG4 activity. Biochemical analyses performed in yeasts (Pérez-Pérez et al., 2014) and Chlamydomonas (Pérez-Pérez et al., 2016) revealed that ATG4 is regulated in these organisms through the formation of a single disulfide bond with a very low redox potential that can be efficiently reduced by thioredoxin. Moreover, it has been shown that both yeast and Chlamydomonas ATG4 proteins can form high molecular mass oligomers under acute oxidizing conditions (Pérez-Pérez et al., 2014; Pérez-Pérez et al., 2016), although the residues responsible for this aggregation have not been identified yet.

Besides ATG4, the formation of ATG8-PE conjugates also depends on ATG7 and ATG3, and a recent study indicated that these two enzymes are also subjected to redox regulation in mammals. *In vitro* and *in vivo* data showed that the oxidation of ATG7 and ATG3 mediates the inhibition of autophagy through the formation of a hetero-disulfide bond that prevents the interaction and lipidation of LC3 (Frudd et al., 2018). The stable covalent complexes between LC3, ATG7 and ATG3 decrease upon autophagy activation when LC3 is transferred to PE. Moreover, the catalytic cys of these two enzymes are prone to oxidation and establish a disulfide bond between ATG7-ATG3 (Frudd et al., 2018). In close agreement with this study, our results showed that ATG3 proteins from Chlamydomonas and yeast are direct targets of ROS and undergo reversible redox PTMs. Our in-depth biochemical analysis demonstrated that the presence of ROS such as H_2_O_2_ provokes the direct oxidation and inhibition of ATG3 (Figures 1–3).

E2-activating ATG3 activity and both proteolytic and deconjugating ATG4 activities should be tightly regulated in order to form ATG8-PE adducts under stress, and Trx, a major cellular thiol-based reductase, might play an important role in this control. We have previously reported that Trx reduces and activates ATG4 in yeast (Pérez-Pérez et al., 2014) and Chlamydomonas (Pérez-Pérez et al., 2016). In the current study, we found that ATG3 requires electron donors to carry out ATG8 lipidation and established this enzyme as a new Trx target (Figure 3). This result is in close agreement with our previous Trx-targeted proteomic analysis in Chlamydomonas where ATG3 was identified as a putative target of cytosolic Trx (Pérez-Pérez et al., 2017). Our in-gel redox titrations indicated that Chlamydomonas and yeast ATG3 enzymes display a very low midpoint redox potential (*E*_m_ −311/-312 mV) (Figure 2), suggesting that these proteins might be regulated by Trx due to its high reducing power (Gelhaye et al., 2004; Yoshida et al., 2018). The ATG3 redox potential values were more negative than those reported for Chlamydomonas and yeast ATG4 proteins (*E*_m_ −278 mV and *E*_m_ −289 mV, respectively) (Pérez-Pérez et al., 2014; Pérez-Pérez et al., 2016). Thus, we hypothesized that ATG4 and ATG3 are coordinately regulated according to the intracellular redox potential. During basal metabolism, the electronegative intracellular redox potential keeps ATG3 and ATG4 mainly reduced and active. The proteolytic ATG4 and conjugating ATG3 activities promote ATG8-PE adduct formation, whereas the delipidating activity of ATG4 prevents excessive ATG8-PE formation, resulting in a basal level of lipidated ATG8 and housekeeping autophagy. By contrast, stress triggers ROS production, turning the redox potential into more electropositive and thus leading to sequential inhibition by oxidation of ATG4 and then ATG3, to facilitate ATG8-PE stability. It is worth mentioning that processing of ATG8 by ATG4 occurs constitutively (Kirisako et al., 2000; Pérez-Pérez et al., 2010), indicating that the ATG4 delipidating activity must be the one highly regulated (Nakatogawa et al., 2012; Yu et al., 2012; Abreu et al., 2017). The redox regulation of these autophagy proteins was accompanied by an upregulation of ATG3 (Figure 6 and Supplemental Figure S7) and ATG4 protein abundance (Pérez-Pérez et al., 2016) upon ROS-mediated activation of autophagy, suggesting that these proteins participate in the cellular response of Chlamydomonas to oxidative stress.

Reversible redox modifications allow proteins to quickly adapt their activity to fluctuating cellular requirements (Choudhury et al., 2017; Wakao and Niyogi, 2021). Indeed, our time-course *in vivo* alkylation shift assays revealed the dynamics of ATG3 redox regulation upon NF-induced carotenoid depletion (Figure 6). Remarkably, the proportion of reduced ATG3 changed in accordance with ROS abundance to ensure ATG8 lipidation and autophagy progression in carotenoid-depleted cells. Norflurazon activates autophagy in Chlamydomonas after 24 h and further stimulates this catabolic process at longer treatments (Figure 6) (Pérez-Pérez et al., 2012a). Our results indicated that norflurazon leads first to oxidation and inactivation of ATG3 at the time of autophagy induction (Figure 6), which is in close agreement with the inactivation of ATG4 reported in NF-treated cells (Pérez-Pérez et al., 2016). However, ATG3 was detected almost exclusively in its reduced and active form under maximal autophagy induction (Figure 6), likely to accomplish ATG8 lipidation and promote autophagy progression. In consonance with these results, alkylation shift assays performed in mammalian cells also revealed increased oxidation of ATG3 in response to H_2_O_2_, which attenuates LC3 lipidation (Frudd et al., 2018).

Our *in vitro* results indicated that ATG3 displays two reversible oxidized isoforms in the presence of ROS inducers, an oxidized monomer and a homodimer (Figure 1), but we could not detect dimeric ATG3 in total extracts from Chlamydomonas using anti-CrATG3 antibodies. Failure to detect oxidized dimers in vivo has been reported for other redox regulated proteins. For instance, the elongation factor EF-Tu from the cyanobacterium Synechocystis is reversibly inhibited by oxidation, forming an intermolecular disulfide bond under oxidizing conditions *in vitro* while no oxidized dimers are detected *in vivo* due to immediate reduction by Trx (Jimbo et al., 2018). Given that ATG3 transiently interacts with ATG7 and ATG8 to transfer ATG8 to PE (Taherbhoy et al., 2011; Kaiser et al., 2012), and ATG3 can also interact with other proteins such as GAPDH (Han et al., 2015), it is plausible that these dynamic interactions prevent the covalent binding between ATG3 molecules and the detection of ATG3 dimers in Chlamydomonas. Otherwise, we cannot rule out that anti-CrATG3 antibodies do not recognize ATG3 oligomeric forms.

Here, we investigated the sensitivity of the catalytic Cys255 of CrATG3 to H_2_O_2_, and the effects of its oxidation on ATG8-PE conjugation activity using a *cell-free* assay with Chlamydomonas total extracts and recombinant proteins. Our results confirmed that ATG3 activity can be monitored in Chlamydomonas whole cell extracts using endogenous ATG8 and recombinant ATG3 protein (Figure 5). This novel assay allowed us to analyze the redox regulation of ATG8 lipidation by ATG3 in a simple way. A similar assay was initially set up to study ATG4 protease activity in Chlamydomonas (Pérez-Pérez et al., 2010) and then in yeasts (Pérez-Pérez et al., 2014) and plants (Laureano-Marín et al., 2020). Therefore, this assay could be exploited for analyzing the regulation of ATG3 activity from other organisms such as yeasts (Figures 5E and 5F) without the limitation of reconstituting the whole ATG8 lipidation system *in vitro* but using the components present in whole cell extracts.

Taken together, our results unravel the molecular mechanisms underlying the redox regulation of ATG3, a main component of the ATG8 lipidation system. The coordinated regulation of different proteins from the ATG8 lipidation system such as ATG3 and ATG4 would connect redox signals and antioxidant systems with the process of autophagy, which functions as a main mechanism to eliminate ROS-damaged components (Pérez-Pérez et al., 2012a; Signorelli et al., 2019).

## MATERIALS AND METHODS

### Strains, media and growth conditions

The *Chlamydomonas reinhardtii* strain used in this study is wild-type 4A+ (CC-4051), which was obtained from the Chlamydomonas Culture Collection. Chlamydomonas cells were grown under continuous illumination (50 μmol photon m^-2^ s^-1^) on an orbital shaker (100 rpm) at 25°C in Tris-acetate phosphate (TAP, pH 7.1) medium as previously described (Harris, 1989). When required, cells growing exponentially (≈10^6^ cells per milliliter) were treated with norflurazon (Sigma-Aldrich, 34364), cerulenin (Sigma-Aldrich, C2389) or metylviologen (Sigma-Alrich, 85617).

The *Saccharomyces cerevisiae* strains used in this study are wild-type (SEY6210) and *atg3* mutant (MT33-1A) and were kindly provided by Dr. Daniel Klionsky (Tsukada and Ohsumi, 1993). Yeast cells were grown in rich medium (YPD; 1% yeast extract [Difco, 10215203], 2% peptone [Difco, 211705], and 2% glucose [wt/vol]) (Sigma-Aldrich, G7021) on an orbital shaker (200 rpm) at 30°C.

### Gene cloning and protein purification

The DNA coding sequence of the *ATG3* genes from *Chlamydomonas reinhardtii* and *Saccharomyces cerevisiae* were synthesized and cloned into pBluescript SK (+) vector by the company GeneCust Europe (genecust.com). *ATG3* coding regions were cloned into the pET28a (+) plasmid (Novagen, 69864-3) for expression of the corresponding His-tagged protein. Chlamydomonas *ATG3* was cloned at *NdeI* and *BamHI* sites of pET28a (+), whereas Saccharomyces *ATG3* was cloned at *BamHI* and *XhoI* sites.

The different Chlamydomonas *ATG3* mutants (*ATG3^C50S^*, *ATG3^C81S^* and *ATG3^C255S^*) and Saccharomyces *ATG3* mutant (*ATG3^C234S^*) were synthesized by GeneCust Europe and cloned into pBluescript SK (+) vector. All *ATG3* mutants were also cloned into the pET28a (+) plasmid at the same restriction sites as wild-type versions. All clones were used to transform *E. coli* BL21 (DE3) strain. Recombinant proteins were expressed by induction at exponential growth phase (OD_600nm_≈0.5) with 0.5 mM isopropyl-β-D-thiogalactopyranoside (Sigma, I6758) for 2.5 h at 37°C. All ATG3 recombinant proteins (wild-type and Cys-to-Ser mutants) were purified by affinity chromatography on a His-Select Nickel Affinity Gel (Sigma, P6611) following manufacturer’s instructions. The wild-type ATG8 from Chlamydomonas was purified as formerly described (Pérez-Pérez et al., 2010). The TRXh1^WT^ protein from Chlamydomonas was purified as described (Goyer et al., 1999). The Chlamydomonas *ATG3* sequence (Cre02.g102350) was obtained from Phytozome (phytozome-next.jgi.doe.gov), whereas the Saccharomyces *ATG3* (YNR007C) sequence was obtained from Yeast Genome Database (yeastgenome.org). All recombinant proteins used in this study are listed in Table S1.

### *In vitro* protein analysis

To investigate the electrophoretic mobility of ATG3 proteins, the typical reaction mixture included 3 μg of ATG3 in 50 mM Tris HCl (pH 7.5) in the absence or presence of reducing [DTT_red_ (Sigma-Aldrich; 43815) or GSH (Sigma-Aldrich; G4251) or oxidizing [DTTox (Sigma-Aldrich; D3511), H_2_O_2_ (Sigma-Aldrich; H1009)] or CuSO4 (Sigma-Aldrich; 451657)] at the indicated concentration and time. For the analysis of thioredoxin activity, 5 μg of Chlamydomonas TRXh1 and 0.5 mM DTT_red_ as electron donor were used at the indicated incubation time. The reaction mixtures were incubated at 25°C for the indicated time and stopped by addition of β-mercaptoethanol (βME)-free loading sample buffer followed by 5 min at 65°C. Proteins were resolved on 12% or 15% SDS-PAGE gels, stained with Coomassie Brilliant Blue (Sigma-Aldrich, 27816) and visualized using a ChemiDoc Imaging System (Bio-Rad, 17001401). When required, protein signals were quantified using the ImageLab software (Bio-Rad) and typically untreated sample was used as reference.

### Redox titration

Redox titration of ATG3 monomerization was performed by incubating the recombinant ATG3 protein from Chlamydomonas or Saccharomyces in 50 mM Tris-HCl (pH 7.5) at defined *E*_h_ values imposed by reduced (DTT_red_) and oxidized (DTTox) DTT in different dithiol/disulfide ratios (with 20 mM as final concentration of total DTT (DTT_red_+DTT_ox_)) as previously described (Pérez-Pérez et al., 2014; Pérez-Pérez et al., 2016). ATG3 samples were equilibrated in redox buffers for 30 min at 25°C and then the reaction was stopped by addition of βME-free loading sample buffer followed by 5 min at 65°C. Then, proteins were separated by 12% SDS-PAGE gels. Proteins were visualized by Coomassie blue staining and ATG3 monomer and dimer were quantified using the ImageLab software (Bio-Rad) and the sample treated with 20 mM DTT_red_ and 0 mM DTTox (*E*_h_= -∞) was used as reference and considered totally reduced. Titration data were fitted to the Nernst equation (n= 2, one component). The midpoint redox potential (*E*_m_) is reported as the mean ± standard deviation of 3 independent experiments. The redox potential values (*E*_h_) are expressed at pH 7.5.

### Protein preparation from Chlamydomonas cells

Chlamydomonas cells from liquid cultures were collected by centrifugation (4000*g* for 4 min), washed in lysis buffer (50 mM Tris-HCl (pH 7.5)), and resuspended in a minimal volume of the same buffer. Cells were lysed by two cycles of slow freezing to −80°C followed by thawing at room temperature. The soluble cell extract was separated from the insoluble fraction by centrifugation (15000*g* for 20 min at 4°C). Proteins were quantified with the Coomassie dye binding method (BioRad, 500-0006) as described by the manufacturer.

### Immunoblot analysis

For immunoblot analyses, total protein extracts (10 to 20 μg) were subjected to 12% or 15% SDS-PAGE and then transferred to 0.45 μm nitrocellulose membranes (GE Healthcare, 10600003). The anti-CrATG3 polyclonal antibody was produced by injecting the recombinant wild-type CrATG3 protein into a rabbit using standard immunization protocols at the Animal Resource facility from the University of Sevilla. Anti-CrATG3 was diluted 1:15000; anti-CrATG8 (Pérez-Pérez et al., 2010), anti-CrATG4 (Pérez-Pérez et al., 2016) and secondary rabbit antibodies were diluted 1:3000, 1:5000, and 1:10000, respectively, in phosphate-buffered saline (PBS) containing 0.1% Tween 20 (Applichem, A4974) and 5% milk powder (Applichem, A0830). The Luminata Crescendo Millipore immunoblotting detection system (Millipore, WBLUR0500) was used to detect proteins with horseradish peroxidase-conjugated anti-rabbit secondary antibodies (Sigma, A6154).

### Cell-free ATG8 lipidation assays in Chlamydomonas and Saccharomyces

Chlamydomonas total cell extracts prepared as described above were incubated at 25°C in the absence or presence of reducing (DTT_red_) and/or oxidizing (H_2_O_2_) agents for the indicated concentration and time, and stopped by addition of βME-free loading sample buffer followed by 5 min at 65°C. Then, proteins were resolved by 15% SDS-PAGE and analyzed by western blot with anti-ATG8 (Pérez-Pérez et al., 2010) as described above.

For ATG8 lipidation assays in yeasts, cells were collected by centrifugation (4000*g* for 4 min), washed in lysis buffer 1 (50 mM Tris-HCl (pH 7.5)), resuspended in lysis buffer 2 (50 mM Tris-HCl (pH 7.5), 1 mM Pefablock (Sigma-Aldrich, 76307), 0.1 % Triton X100 (Sigma-Aldrich, T9284)) and vortexed 10 times for 30 s each time in the presence of glass beads (Sigma-Aldrich, G8772). Total cell extract was separated by centrifugation (15000*g* for 20 min at 4°C). Proteins were incubated at 25°C with recombinant Chlamydomonas His-tagged ATG8 in the absence or presence of DTT_red_/H_2_O_2_ and ATG8 lipidation was analyzed by western blot analysis with Chlamydomonas anti-ATG8 antibody (Pérez-Pérez et al., 2010) as described above.

### Redox western blots

Cys-alkylation assays were performed using N-ethylmaleimide (NEM) (Sigma-Aldrich, E1271) and Methyl-PEG-Maleimide Reagent (MM(PEG)_24_) (Thermo Fisher, 22713). NEM was added to Chlamydomonas cultures growing as indicated in each case to a final concentration of 10 mM. After 15 min of NEM incubation, cells were collected by centrifugation (4000*g* for 4 min), washed in alkylating buffer (50 mM Tris-HCl (pH 7.5), 50 mM NaCl and 10 mM NEM), and resuspended in a minimal volume of the same buffer. Cells were lysed by two cycles of slow freezing to −80°C followed by thawing at room temperature. The soluble cell extract was separated from the insoluble fraction by centrifugation (15000*g* for 20 min at 4°C), and incubated with 10% (w/v) of trichloroacetic acid (TCA) (Sigma-Aldrich, T6399) for 30 min on ice. Then, samples were centrifugated (15000*g* for 20 min at 4°C), and precipitates were washed twice with ice-cold acetone and resuspended in a SDS-containing buffer (50 mM Tris-HCl (pH 7.5), 50 mM NaCl and 2% (w/v) SDS). Next, samples were incubated with 100 mM DTT_red_ for 60 min on ice. Finally, samples were again treated with TCA and ice-cold acetone and precipitates were resuspended in a new buffer (50 mM Tris-HCl (pH 8.0), 50 mM NaCl, 2% (w/v) SDS, 7.5% (v/v) glycerol and 0.01% bromo-phenol blue) in the absence or presence of 10 mM MM(PEG)_24_.

## Acknowledgments and funding

We would like to thank Dr. Christophe Marchand for sharing Chlamydomonas TRXh1 protein and Professor D. Klionsky for kindly providing yeast wild-type (SEY6210) and *atg3* (MT33-1A) mutant strains. We would like to specially thank Dr. José L. Crespo for discussion, data analysis and the critical reading of the manuscript. This work was supported in part by the Ministerio de Economía y Competitividad (grant BIO-2015-74432-JIN to M.E.P.-P.) and Ministerio de Ciencia e Innovación (grant PID2019-110080GB-100 to M.E.P.-P.).

## Author contributions

M.E.P.-P. designed research and wrote the manuscript with the input of all authors. M.J.M.-P. and M.E.P.-P. performed research. All authors reviewed and edited the manuscript. There is no conflict of interest.

