## Supplemental Material for "Redox-mediated activation of ATG3 promotes ATG8 lipidation and autophagy progression in Chlamydomonas"

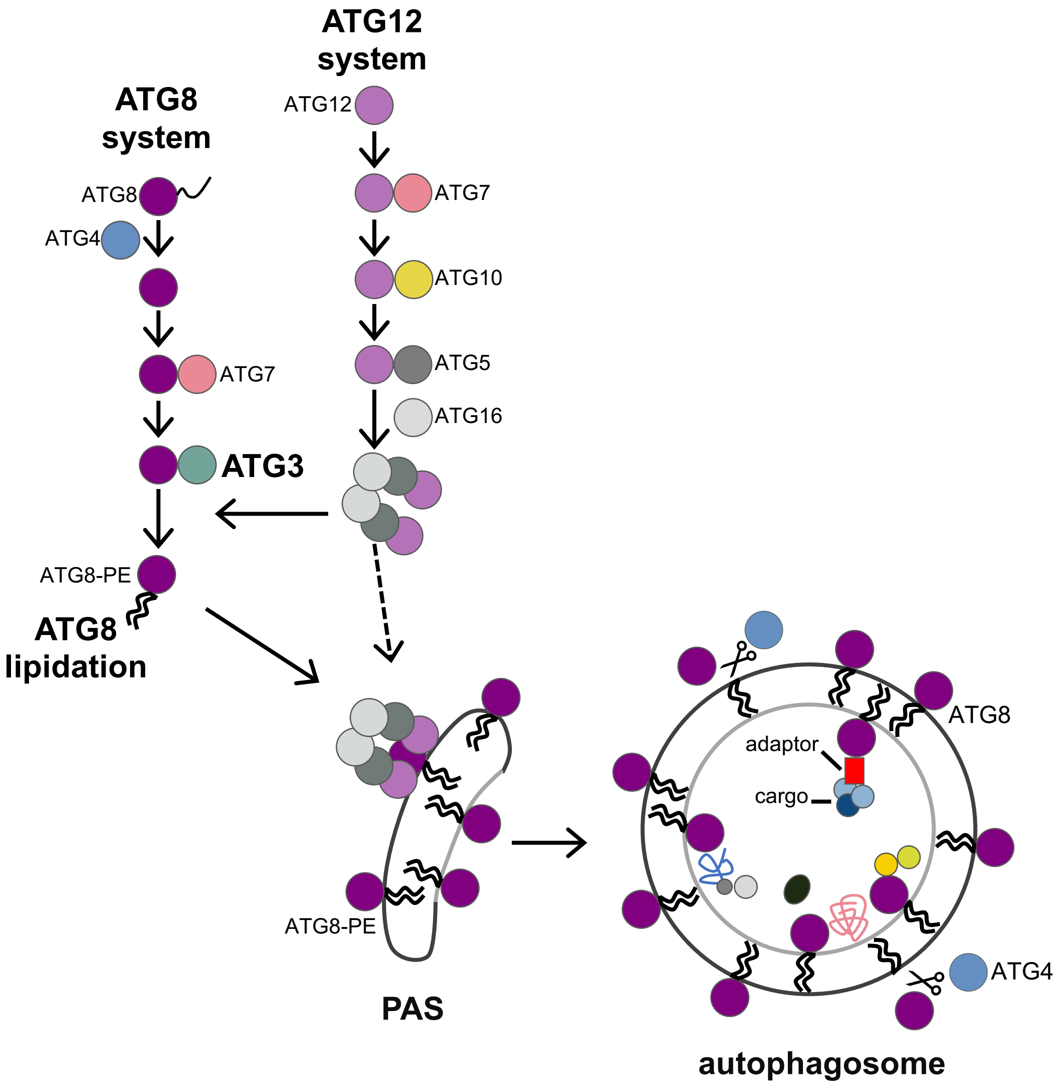


**Figure S1. The ATG8 and ATG12 ubiquitin-like systems in autophagy.** After autophagy initiation, the components of ATG8 (ATG3, ATG4, ATG7 and ATG8 proteins) and ATG12 (ATG5, ATG7, ATG10, ATG12 and ATG16 proteins) systems are recruited to the preautophagosomal structure or PAS. All of these proteins act coordinately and sequentially to promote autophagosome biogenesis. The conjugation of ATG8 to the membrane phospholipid phosphatidylethanolamine (PE) is a key step in vesicle expansion and autophagosome formation. For ATG8 lipidation, first, nascent ATG8 is cleaved at a highly conserved Gly at its C-terminus by the Cys-protease ATG4. The C-terminal Gly is then activated by the E1 enzyme ATG7, with ATP consumption, and transferred to the E2 enzyme ATG3. Finally, ATG8 is bound to the headgroup of PE, and the ATG8-PE adduct is formed. The ATG5-ATG12-ATG16 protein complex acts as an E3 enzyme to stimulate the binding of ATG8 to PE. Besides processing ATG8 precursor, ATG4 presents a deconjugating or delipidating activity to cleave ATG8-PE and release free ATG8 from autophagosome membranes for recycling. The lipidation of ATG8 also plays an important role in cargo recognition since ATG8-PE interacts specifically with the intracellular material that will be engulfed into the autophagosome directly or through adaptors.

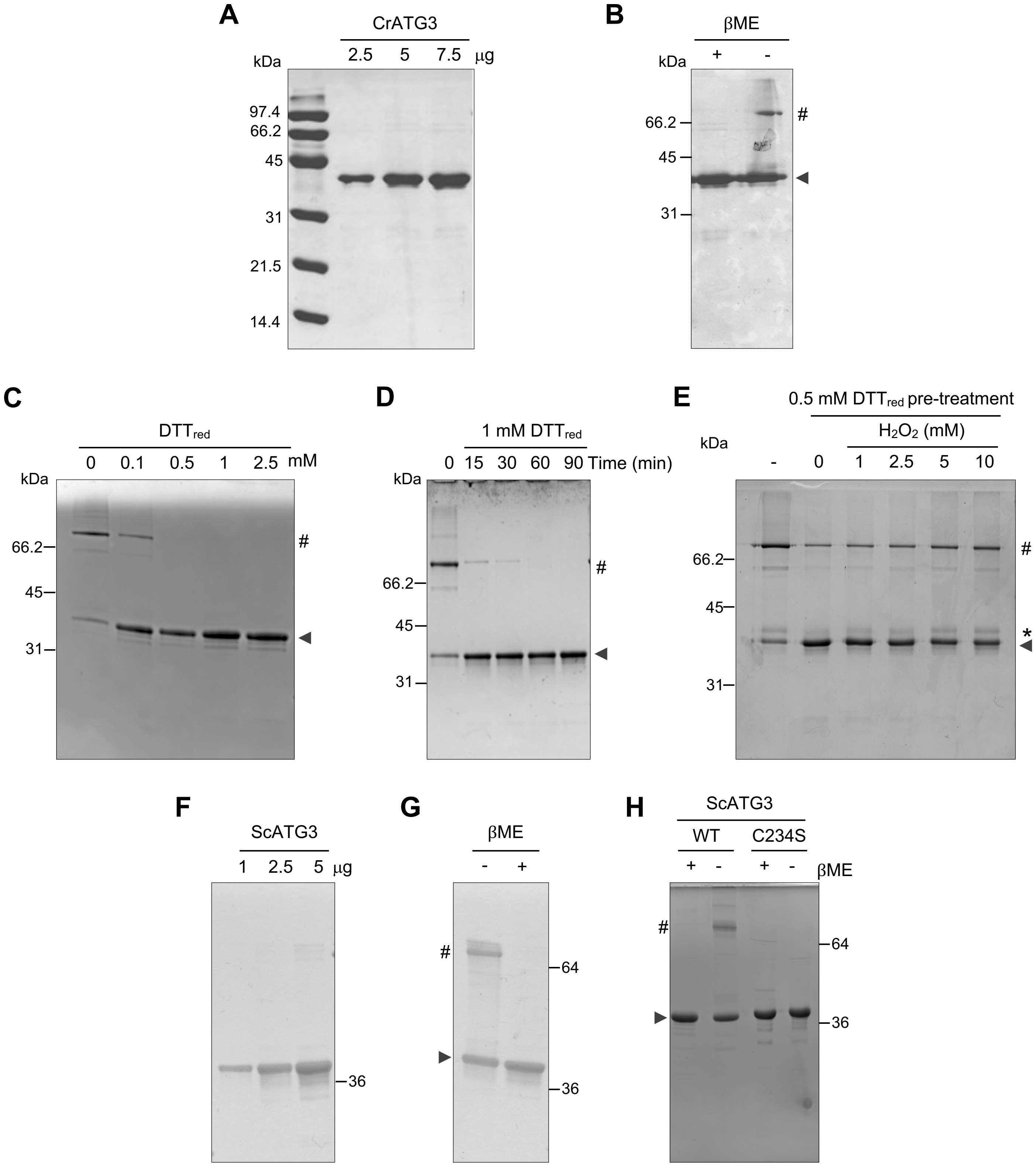


**Figure S2. His-tagged recombinant ATG3 proteins from Chlamydomonas (CrATG3) and Saccharomyces (ScATG3) purified in *E. coli*. A.** Increasing concentrations (2.5, 5 and 7.5 μg) of CrATG3 in a β-mercaptoethanol (βME)-containing loading buffer subjected to SDS-PAGE gel. **B.** Electrophoretic mobility of CrATG3 in the absence (-) or presence (+) of the reducer βME in a non-reducing SDS-PAGE gel. **C.** CrATG3 treated with increasing concentrations of DTT$_{red}$ (0, 0.5, 1 and 2.5 mM) for 30 min. **D.** CrATG3 treated with 1 mM DTT$_{red}$ along time (0, 15, 30, 60 and 90 min). **E.** CrATG3 in the presence of increasing concentrations of $H_2O_2$ (0, 1, 2.5 and 5 mM) for 30 min after pretreatment with DTT$_{red}$ (0.5 mM, 60 min). The molecular mass marker (kDa) is shown on the left. The different CrATG3 isoforms are indicated with an arrow (reduced monomeric ATG3), an asterisk (*) (oxidized monomeric ATG3) and a hash (#) (dimeric ATG3).

**A**

| Organism | ATG3 Chlamydomonas (% Identity) | Amino acid number (Cys number) |
|---|---|---|
| ***Chlamydomonas reinhardtii*** | **100** | **306 (3)** |
| *Volvox carteri* | 69.3 | 332 (3) |
| *Chlorella variabilis* | 53.1 | 330 (3) |
| *Ostreococcus lucimarinus* | 43.6 | 290 (3) |
| *Phaeodactylum tricornutum* | 38.7 | 302 (4) |
| *Arabidopsis thaliana* | 50.6 | 313 (3) |
| *Zea mays* | 50.3 | 311 (3) |
| *Oryza sativa* | 49.5 | 316 (3) |
| *Schyzosaccharomyces pombe* | 34.2 | 275 (4) |
| ***Saccharomyces cerevisiae*** | **32** | **310 (4)** |
| *Drosophila melanogaster* | 41.2 | 330 (5) |
| *Homo sapiens* | 36.9 | 317 (8) |

**B**

```
CrATG3      MSNLRHTLHTLFKQTVETVTPPLTKSQFEEKRVLTPDEFVAAGDYLVHACPTWSWEGGDP  60
ScATG3      ------MIRSTLSSWREYLTPITHKSTFLTTGQITPEEFVQAGDYLCHMFPTWKWNEESS  54
                   ::: :..  * :**   ** *  .  :**:*** ***** *  ***.*:  .

CrATG3      -KKRRTYFPPNKQFLVTRNVPCLKRATELEGYNPNSEFDVGGG-EGEDAWVATHSNPAAA  118
ScATG3      DISYRDFLPKNKQFLIIRKVPCDKRAEQCVEVEGPDVIMKGFAEDGDED-----------  103
              . * ::* *****: *:*** *** :    :  . :  * . :*::

CrATG3      SGSAGKGEVPSIDGAGAGGSGGAGAAGGNKDDDIPDITD----LELNEADDEAAAPSGRP  174
ScATG3      -------DVLEYIGSET-EHVQSTPAGGTKDSSIDDIDELIQDMEIKEEDEN---DDTEE  152
                   :* .  *: :     :  ***.**..* ** :    :*::* *::    . .

CrATG3      YLRAEEPADNIMRTRTYDLYITYDQYYQVPRFWLVGHDESRKPLLPQQVMEDVSEEHARK  234
ScATG3      FNAKGGLAKDMAQERYYDLYIAYSTSYRVPKMYIVGFNSNGSPLSPEQMFEDISADYRTK  212
            :      *.:: : * *****:*.  *:**:::**.:..  .** *:*::**:*  ::  *

CrATG3      TITVDPHPHLA-GLSAASIHPCRHADVMKKLVDNLLE-----------------------  270
ScATG3      TATIEKLPFYKNSVLSVSIHPCKHANVMKILLDKVRVVRQRRRKELQEEQELDGVGDWED  272
            * *::  *.   .: :.*****:**:*** *:*::

CrATG3      ----AGREFKVEQYLVLFLKFIASVVPTIQYDYTMSVGGE* 306
ScATG3      LQDDIDDSLRVDQYLIVFLKFITSVTPSIQHDYTMEGW--- 310
                 . .::*:***::*****:**.*:**:****.
```

**Figure S3. Comparison of ATG3 enzymes from different organisms. A.** Table showing the identity (%) of ATG3 proteins from different species (*Chlamydomonas reinhardtii; Volvox carteri; Chlorella variabilis; Ostreococcus lucimarinus; Phaeodactylum tricornutum; Arabidopsis thaliana; Zea mays; Oryza sativa; Schyzosaccharomyces pombe; Saccharomyces cerevisiae; Drosophila melanogaster* and *Homo sapiens*). The identity was obtained by alignment of the two sequences using Pairwise Sequence Alignment EMBOSS Needle. The identity (%), the amino acid and cysteine residues numbers (in parenthesis) are indicated. The two organisms used in this study (*Chlamydomonas reinhardtii* and *Saccharomyces cerevisiae*) are highlighted in bold. The accession numbers and the species abbreviations are indicated. CrATG3: *Chlamydomonas reinhardtii* (EDP07491); VcATG3: *Volvox carteri* (EFJ46364); CvATG3: *Chlorella variabilis* (EFN54110); OlATG3: *Ostreococcus lucimarinus* (ABO96836); Pt: *Phaeodactylum tricornutum* (EEC51122); At: *Arabidopsis thaliana* (OAO94560); ZmATG3: *Zea mays* (ACJ72033); OsATG3: *Oryza sativa* (EEE54060.1); SpATG3: *Schyzosaccharomyces pombe* (CAA17786); *Saccharomyces cerevisiae* (KZV08635); DmATG3: *Drosohpila melanogaster* (NP_649059); *Homo sapiens* (AAH02830). ATG3 sequences from all organisms were obtained from NCBI (https://www.ncbi.nlm.nih.gov/pubmed). **B.** Amino acid sequence alignment of CrATG3 and ScATG3. The proteins were aligned with the Clustal Omega software. All Cys residues are highlighted in red. The identical residues in both sequences are shown in bold.

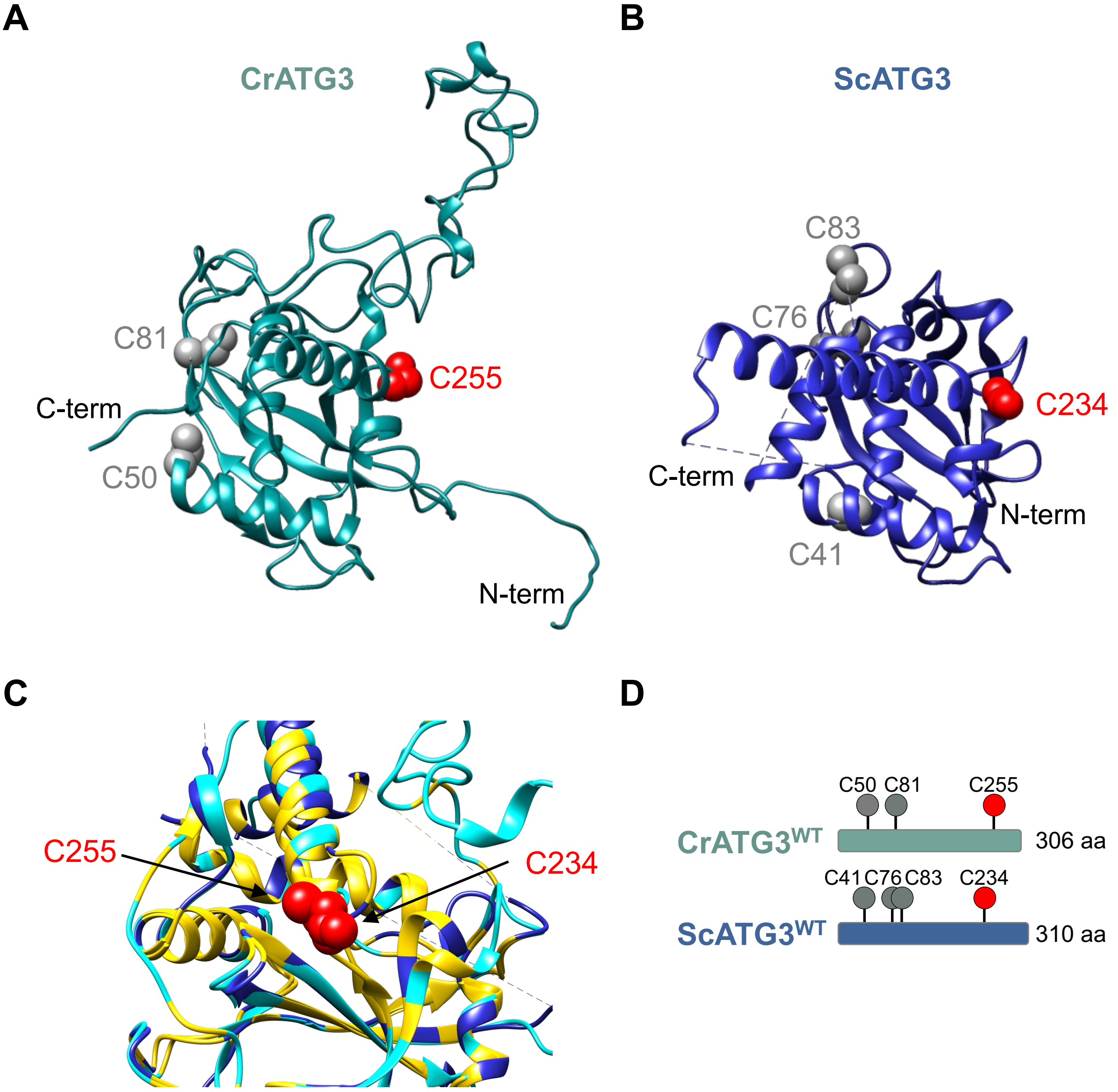


**Figure S4. Predicted structure of ATG3 enzyme form Chlamydomonas based on the structure of Saccharomyces ATG3 (PDB ID: 2DYT). A-B.** Predicted structure of CrATG3 *(A)* and ScATG3 *(B)* showing all Cys of each protein and the N- and C-terminal regions. **C.** Enlarged overlapping of both predicted structures showing the catalytic Cys (Cys255 in CrATG3 and Cys234 in Saccharomyces) and the highly conserved area surrounding the catalytic Cys. CrATG3 is shown in turquoise, ScATG3 in blue, and the overlapping structure of both ATG3 is shown in yellow. **D.** Schematic representation of CrATG3 and ScATG3 showing the amino acid number. All Cys residues present in each ATG3 sequence are depicted: the highly conserved catalytic Cys (Cys255 in Chlamydomonas and Cys234 in Saccharomyces) is highlighted as a red ball, whereas the rest of Cys (Cys50 and Cys81 in Chlamydomonas; Cys71, Cys76 and Cys83 in Saccharomyces) are indicated as grey balls.

**A**

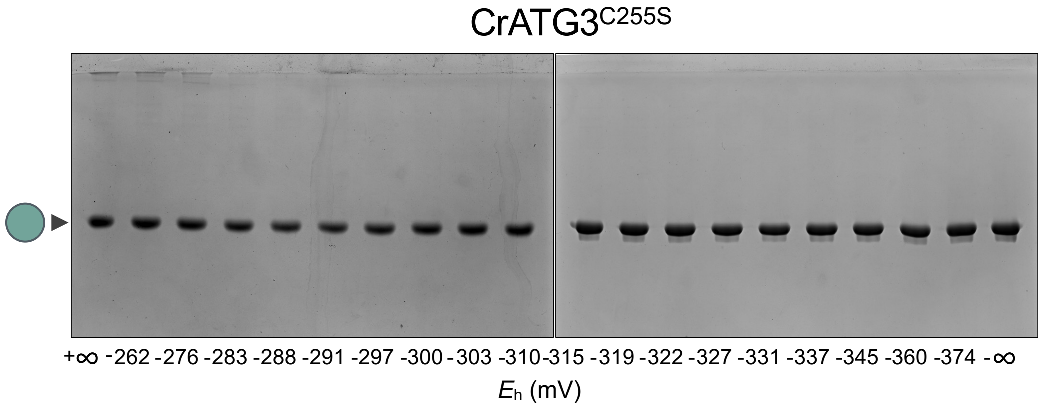


**Figure S5. Redox titration of Chlamydomonas catalytic Cys-to-Ser CrATG3 mutant protein monomerization. A.** SDS-PAGE of CrATG3$^{C255S}$ lacking of Cys255. The CrATG3$^{C255S}$ monomer proportion was monitored after incubation at indicated $E_h$ poised by 20 mM DTT in various dithiol/disulfide ratios (indicated at pH 7.5). All samples were resolved by non-reducing SDS-PAGE gels and then visualized by Coomassie Brilliant Blue. The -∞ and +∞ samples correspond to incubation with only $DTT_{red}$ or $DTT_{ox}$, respectively. The monomeric CrATG3$^{C255S}$ is indicated with a single green ball.

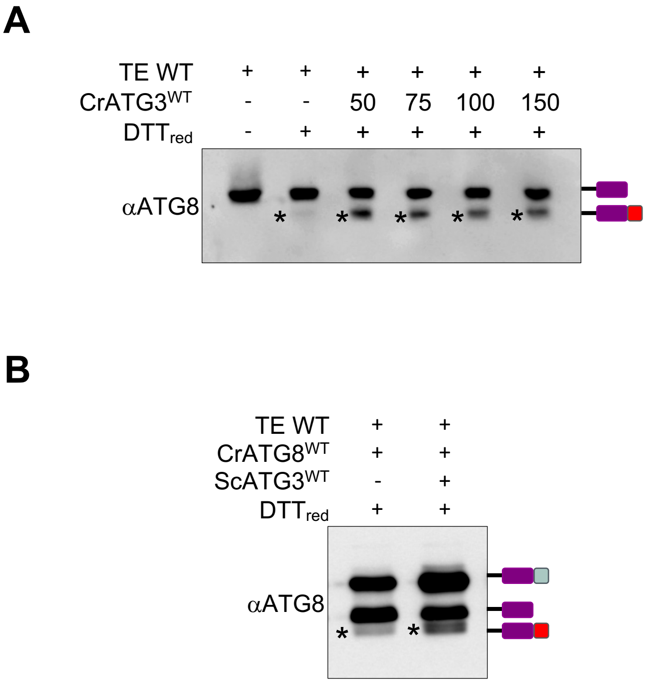


**Figure S6. Reduced ATG3 mediates Chlamydomonas ATG8 lipidation in cell-free assays. A.** Total extracts from Chlamydomonas WT were untreated (-) or treated (+) with $DTT_{red}$ in the absence (-) or presence of increasing concentrations of recombinant $CrATG3^{WT}$ for 60 min at 25ºC. **B.** Total extracts from Saccharomyces WT in a $DTT_{red}$-containing buffer were incubated with recombinant $CrATG8^{WT}$ in the absence (-) or presence (+) of recombinant $ScATG3^{WT}$ for 60 min at 25ºC. After the assay, proteins were subjected to SDS-PAGE gel and then to western blot analysis with Chlamydomonas anti-ATG8 antibody (αATG8). The different protein versions of CrATG8 (unprocessed, processed and lipidated CrATG8) are indicated on the right. CrATG8-PE is highlighted with an asterisk (on the gel).

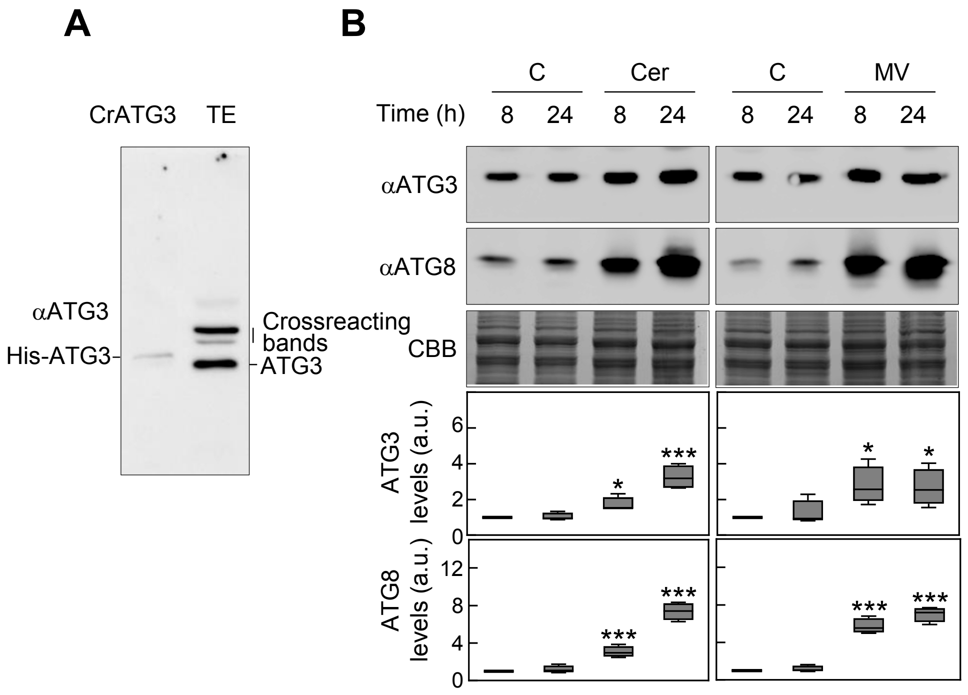


**Figure S7. CrATG3 protein levels are upregulated under autophagy-activating conditions in Chlamydomonas. A.** His-tagged ATG3 (5 ng) and total extracts (20 μg) from Chlamydomonas growing exponentially under optimal conditions were subjected to western blot with Chlamydomonas ATG3 antibodies. **B.** Immunoblot analysis of Chlamydomonas ATG3 (upper panel) and ATG8 (middle panel) in the presence of 10 μM cerulenin (Cer) or 1 μM methyl viologen (MV) for 8 and 24 h. Untreated cells (C) were used as control. Coomassie brilliant blue-stained (CBB) gels were used as protein loading control. Quantification of ATG3 levels (upper box plot) and ATG8 levels (lower box plot) from at least four biological replicates are shown. Asterisks represent significant differences according to two-tailed Student's t test, ***$P < 0.001$ and *$P < 0.05$

**Table S1.** Proteins used in this study.

| Name | Characteristics | Reference |
|---|---|---|
| **CrATG8$^{WT}$** | His-tag WT ATG8 from Chlamydomonas | 1 |
| **CrATG3$^{WT}$** | His-tag WT ATG3 from Chlamydomonas | This study |
| **CrATG3$^{C50S}$** | His-tag ATG3 where Cys50 was replaced by Ser from Chlamydomonas | This study |
| **CrATG3$^{C81S}$** | His-tag ATG3 where Cys81 was replaced by Ser from Chlamydomonas | This study |
| **CrATG3$^{C255S}$** | His-tag ATG3 where the catalytic Cys255 was replaced by Ser from Chlamydomonas | This study |
| **ScATG3$^{WT}$** | His-tag WT ATG3 from Saccharomyces | This study |
| **ScATG4$^{C234S}$** | His-tag ATG3 where Cys234 was replaced by Ser from Saccharomyces | This study |
| **CrTRXh1** | His-tag WT TRXh1 from Chlamydomonas | 2 |

**Cr:** *Chlamydomonas reinhardtii*; **Sc:** *Saccharomyces cerevisiae*.

1. Pérez-Pérez ME, Florencio FJ, Crespo JL (2010). Plant Physiol, 152(4):1874-1888.
2. Goyer A, Decottignies P, Lemaire S, Ruelland E, Issakidis-Bourguet E, Jacquot J-P, Miginiac-Maslow M (1999). FEBS Lett., 444(2-3):165-169.

**Table S2. Statistical analysis from main figures.**

| Figure | Experimental condition | Statistical analysis | n | *P* value |
|---|---|---|---|---|
| 6A | αATG3 (8 h control vs 8 h NF) | Two-tailed Student´s *t* test | 4 | Not significant |
| 6A | αATG3 (24 h control vs 24 h NF) | Two-tailed Student´s *t* test | 4 | *P*=0.00068 *** |
| 6A | αATG8 (8 h control vs 8 h NF) | Two-tailed Student´s *t* test | 4 | Not significant |
| 6A | αATG8 (24 h control vs 24 h NF) | Two-tailed Student´s *t* test | 4 | *P*<0.00001 *** |
| 6C | ATG3 Reduced form (%) (0 h vs 8 h NF) | One-way ANOVA and Bonferroni´s test | 3 | Not significant |
| 6C | ATG3 Reduced form (%) (0 h vs 24 h NF) | One-way ANOVA and Bonferroni´s test | 3 | *P*=0.004 ** |
| 6C | ATG3 Reduced form (%) (0 h vs 48 h NF) | One-way ANOVA and Bonferroni´s test | 3 | *P*=0.015 * |
| S7B | αATG3 (8 h control vs 8 h Cer) | Two-tailed Student´s *t* test | 4 | *P*=0.01428 * |
| S7B | αATG3 (24 h control vs 24 h Cer) | Two-tailed Student´s *t* test | 4 | *P*=0.00075 *** |
| S7B | αATG3 (8 h control vs 8 h MV) | Two-tailed Student´s *t* test | 4 | *P*=0.01124 * |
| S7B | αATG3 (24 h control vs 24 h MV) | Two-tailed Student´s *t* test | 4 | *P*=0.02704 * |
| S7B | αATG8 (8 h control vs 8 h Cer) | Two-tailed Student´s *t* test | 4 | *P*=0.00038 *** |
| S7B | αATG8 (24 h control vs 24 h Cer) | Two-tailed Student´s *t* test | 4 | *P*<0.00001 *** |
| S7B | αATG8 (8 h control vs 8 h MV) | Two-tailed Student´s *t* test | 4 | *P*=0.00003 *** |
| S7B | αATG8 (24 h control vs 24 h MV) | Two-tailed Student´s *t* test | 4 | *P*<0.00001 *** |

*** *P*<0.001; ** *P*<0.01; **P*<0.05; Not significant ≥0.05
